# Precise temporal regulation of post-transcriptional repressors is required for an orderly *Drosophila* maternal-to-zygotic transition

**DOI:** 10.1101/862490

**Authors:** Wen Xi Cao, Sarah Kabelitz, Meera Gupta, Eyan Yeung, Sichun Lin, Christiane Rammelt, Christian Ihling, Filip Pekovic, Timothy C. H. Low, Najeeb U. Siddiqui, Matthew H. K. Cheng, Stephane Angers, Craig A. Smibert, Martin Wühr, Elmar Wahle, Howard D. Lipshitz

## Abstract

In animal embryos the maternal-to-zygotic transition (MZT) hands developmental control from maternal to zygotic gene products. We show that the maternal proteome represents over half of the protein coding capacity of the *Drosophila melanogaster* genome and that 2% of this proteome is rapidly degraded during the MZT. Cleared proteins include the post-transcriptional repressors Cup, Trailer hitch (TRAL), Maternal expression at 31B (ME31B), and Smaug (SMG). While the ubiquitin-proteasome system is necessary for clearance of all four repressors, distinct E3 ligase complexes target them: the C-terminal to Lis1 Homology (CTLH) complex targets Cup, TRAL and ME31B for degradation early in the MZT; the Skp/Cullin/F-box-containing (SCF) complex targets SMG at the end of the MZT. Deleting the C-terminal 233 amino acids of SMG makes the protein immune to degradation. We show that artificially persistent SMG downregulates the zygotic re-expression of mRNAs whose maternal contribution is cleared by SMG. Thus, clearance of SMG permits an orderly MZT.

## INTRODUCTION

Embryonic development in all animals begins with the maternal-to-zygotic transition (MZT) (Tadros and Lipshitz, 2009; Vastenhouw et al., 2019). The MZT can be divided into two phases: Initially, maternally supplied RNAs and proteins direct embryonic development; subsequently, activation of transcription from the zygotic genome, a process termed ‘zygotic genome activation’ (ZGA), transfers developmental control from the mother’s genome to that of the embryo. During the first phase, post-transcriptional regulation of maternal transcripts and post-translational regulation of maternal proteins predominate. The former is coordinated by RNA-binding proteins (RBPs), which regulate the translation, stability and localization of the maternal transcripts. A large proportion of maternal mRNA species is degraded in a highly coordinated manner during the MZT (Aanes et al., 2014; De Renzis et al., 2007; Laver et al., 2015; Stoeckius et al., 2014; Svoboda et al., 2015; Tadros et al., 2007; Thomsen et al., 2010). Transcriptome-wide changes in the translational status of mRNAs have also been described (Chen et al., 2014; Eichhorn et al., 2016; Rissland et al., 2017; Subtelny et al., 2014; Wang et al., 2017; Winata et al., 2018) and global changes in the proteome have been documented (Baltz et al., 2012; Becker et al., 2018; Casas-Vila et al., 2017; Fabre et al., 2016; Gouw et al., 2009; Kronja et al., 2014a; Peshkin et al., 2015; Stoeckius et al., 2014; Sysoev et al., 2016).

Relevant to the changes in the proteome is the ubiquitin-proteasome system, which is a highly conserved and widespread pathway for specific targeting of proteins for degradation (Komander and Rape, 2012; Ravid and Hochstrasser, 2008). This is accomplished through the E1-E2-E3 enzyme ubiquitination cascade, with the E3 ubiquitin ligase acting as the substrate-specificity factor, which transfers ubiquitin from an E2 ubiquitin-conjugating enzyme to specific target proteins (Pickart, 2001; Zheng and Shabek, 2017). Regulation of protein stability by the ubiquitin-proteasome system during the MZT has been noted in several studies. For example, MG132-directed inhibition of maternal protein degradation in mouse early zygotes delays ZGA (Higuchi et al., 2018). Also in mouse, loss of an E3 ubiquitin ligase, RNF114, prevents development beyond the two-cell stage (Yang et al., 2017). RNF114-directed ubiquitination and clearance of TAB1 permits NF-κB pathway activation, although why this is necessary for the MZT is not known. In *C. elegans*, E3-ligase-directed clearance of the RBPs, OMA-1 and OMA-2, in the early embryo is crucial for the temporal coordination of ZGA (Du et al., 2015; Guven-Ozkan et al., 2008; Kisielnicka et al., 2018; Shirayama et al., 2006; Tsukamoto et al., 2017).

In *Drosophila melanogaster*, Smaug (SMG), a multifunctional RBP, is essential for both maternal mRNA degradation and ZGA (Benoit et al., 2009). SMG protein accumulates rapidly at the onset of embryogenesis, when the Pan gu (PNG) kinase complex abrogates translational repression of the *smg* and *Cyclin B* (as well as many other) mRNAs (Kronja et al., 2014b; Tadros et al., 2007; Vardy and Orr-Weaver, 2007). SMG binds target mRNAs through a stem-loop structure known as the Smaug recognition element or ‘SRE’ (Aviv et al., 2006; Aviv et al., 2003). Through these elements SMG induces degradation and/or represses the translation of a large subset of the maternal transcripts (Chen et al., 2014; Semotok et al., 2005; Semotok et al., 2008; Tadros et al., 2007; Zaessinger et al., 2006). SMG down-regulates target mRNA expression through the recruitment of proteins that influence how these mRNAs interact with the mRNA decay and translation machineries. For example, SMG recruits the CCR4-NOT deadenylase complex to induce transcript degradation (Semotok et al., 2005) and the *Drosophila* miRNA Argonaute (AGO), AGO1, to repress mRNA translation (Pinder and Smibert, 2013). SMG acts in a complex with additional translational repressors, including the eIF-4E-binding protein, Cup; the DEAD-box helicases, ME31B and Belle; and the FDF-domain protein, Trailer hitch (TRAL) (Götze et al., 2017; Jeske et al., 2011; Nakamura et al., 2001; Nakamura et al., 2004; Nelson et al., 2004; Wilhelm et al., 2003; Wilhelm et al., 2000). PNG is required for the degradation of the Cup, TRAL and ME31B repressors in early embryos (Wang et al., 2017) and at least one of these, TRAL, may be a direct substrate of PNG (Hara et al., 2018). SMG protein itself is rapidly degraded at the end of the MZT (Benoit et al., 2009; Siddiqui et al., 2012) but the mechanisms and functions of SMG clearance are unknown.

Leveraging the increasing sensitivity and measurement quality of multiplexed proteomics (Pappireddi et al., 2019; Sonnett et al., 2018b), we present here a quantification of the developmental proteome of the *Drosophila* embryo with a focus on the MZT. We show that the embryonic proteome represents over half of the protein-coding capacity of the genome. We reveal a distinct cluster of proteins, comprising 2% of the embryonic proteome, which are highly expressed at the beginning of the MZT but are then rapidly degraded. This cluster includes SMG, Cup, TRAL and ME31B. Focusing on these four repressors, we find that they can be subdivided into two classes: Cup, TRAL and ME31B are degraded rapidly early in the MZT whereas SMG is degraded later, towards the end of the MZT. We identify two distinct E3 ubiquitin ligase complexes that target the classes through the ubiquitin proteasome: the CTLH complex, which is homologous to the yeast Gid-complex (Francis et al., 2013; Liu and Pfirrmann, 2019), targets Cup, TRAL and ME31B; the SCF complex (Ho et al., 2006) targets SMG. We then engineer a stable version of the SMG protein that does not undergo degradation at the end of the MZT and show that persistent SMG downregulates zygotic re-expression of a subset of its maternal targets. Thus, clearance of SMG is necessary for this aspect of the MZT.

## RESULTS

### Definition and dynamics of the embryonic proteome

We performed quantitative complement tandem mass tag (TMTc+) mass spectrometry (Sonnett et al., 2018b) on the proteome of embryos at three stages spanning the MZT, as well as at two later stages: (1) early syncytial blastoderm, prior to zygotic genome activation (ZGA); (2) nuclear cycle 14 (NC14) during blastoderm cellularization, after high-level ZGA; (3) germ-band extension, representing the end of the MZT; (4) germ-band retraction, representing mid-embryogenesis; and (5) tracheal filling, shortly before the end of embryogenesis (Figure 1A). We chose TMTc+ for this analysis because, in comparison with label-free proteomics, it generates data with higher measurement precision and overcomes the ‘missing value’ problem (Pappireddi et al., 2019). Furthermore, TMTc+ eliminates ratio-compression, which is a common problem in other multiplexed proteomics approaches (Pappireddi et al., 2019; Sonnett et al., 2018b; Ting et al., 2011). In total we detected 7564 unique proteins encoded by 7317 genes (Supplemental file 1), representing 53% of the protein-coding genome (7317/13918 per FlyBase Release 6.03) (Matthews et al., 2015). This represents 40% more proteins than previously reported to be expressed during embryogenesis with label-free proteomics (Casas-Vila et al., 2017). Our quantitative proteomics data provide unprecedented depth and measurement quality and should serve as a useful resource for the community.

**Figure 1.**
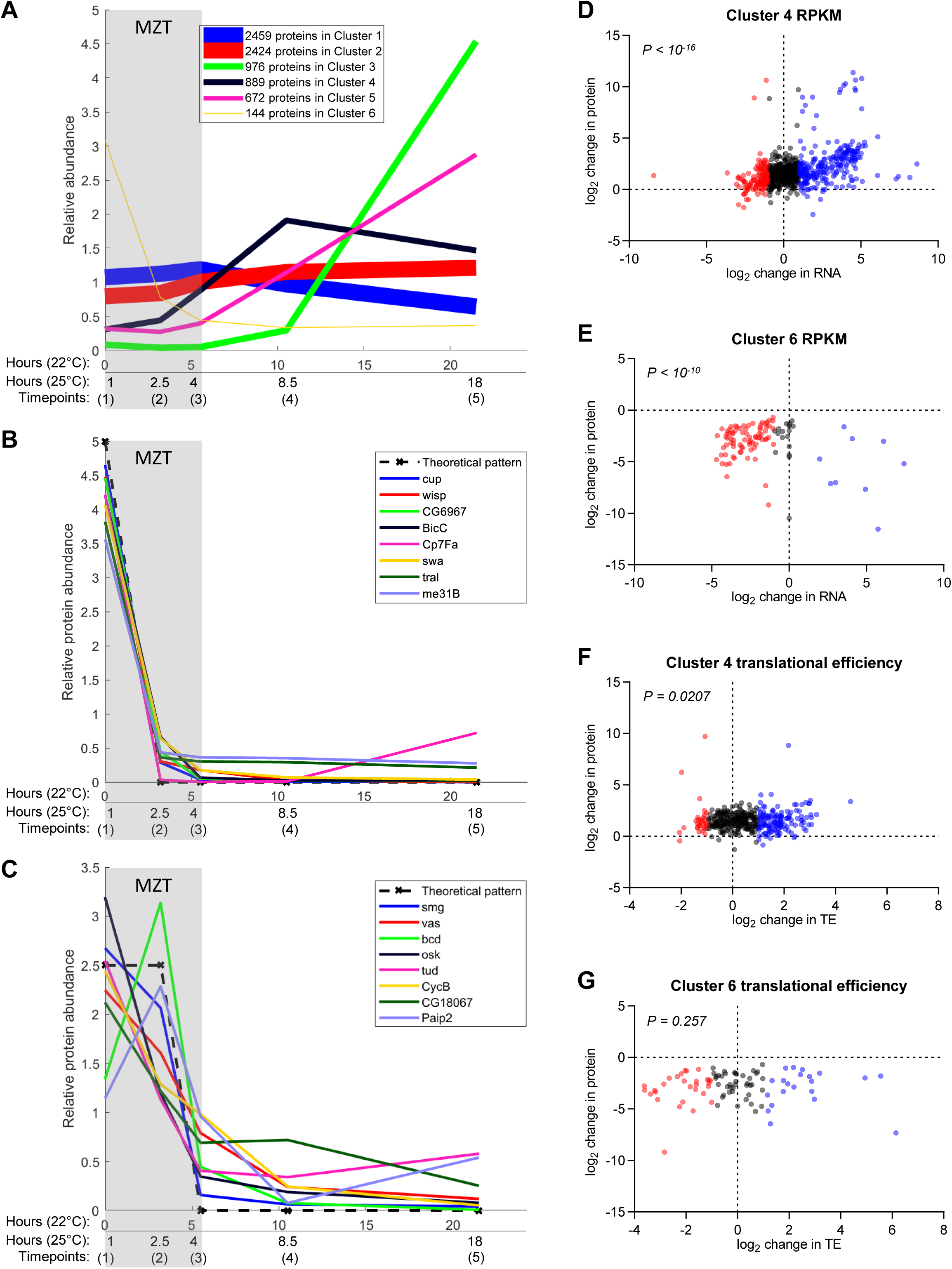
The *Drosophila* proteome is dynamic during embryogenesis. **A.** k-means clustering (k=6) for 7564 quantified proteins through embryogenesis. Embryos were aged to five developmental timepoints at 22°C. The equivalent developmental times at 25°C for each sample are indicated on the x-axis. The first three samples cover the MZT (grey box). **B-C.** Degradation dynamics of RNA-binding proteins in Cluster 6 can be closely modelled by two distinct “theoretical patterns” during the MZT. **B.** Those degraded before Timepoint 2 (nuclear cycle 14) include Cup, TRAL and ME31B. **C.** Those degraded after Timepoint 2, towards the end of the MZT, include SMG. **D-E.** Scatter plot of change in RNA expression (RPKM, 3-4h / 0-1h) (Eichhorn et al., 2016) versus change in protein expression (Timepoint 3 / Timepoint 1) for dynamic proteins during the MZT**. D.** Proteins that increase in expression during the MZT (Cluster 4) significantly correspond with ≥ 2-fold increase in their transcript expression (*P* < *10*^*-17*^); **E.** Proteins that decrease in expression (Cluster 6) significantly correspond with ≥ 2-fold decrease in their transcript expression during the MZT (*P* < *10*^*-11*^). **F-G.** Scatter plot of change in translational efficiency (TE, 3-4h / 0-1h) (Eichhorn et al., 2016) versus change in protein expression (Timepoint 3 / Timepoint 1) for dynamic proteins during the MZT**. F.** Cluster 4 proteins significantly correspond with ≥ 2-fold increase in TE of their transcripts (*P = 0.0207*); **G.** Cluster 6 proteins are not associated with significant changes in TE of their transcripts (*P = 0.257*). Fisher’s exact test performed for **D-G**.

K-means clustering of relative protein expression revealed several classes of proteins with distinct expression dynamics during and after the MZT (Figure 1A). A majority (65%) of proteins was present throughout the MZT and later embryogenesis undergoing less than two-fold increases or decreases in relative levels over the time-course (Clusters 1 and 2: 2459 and 2424 proteins, respectively). These likely represent stable maternally encoded proteins, which are predominantly already deposited in the egg, similar to what has been observed in frog embryos (Peshkin et al., 2015). Alternatively, rates of synthesis and degradation could be similar thereby resulting in relatively constant levels of these proteins (Kronja et al., 2014a). Proteins in Clusters 3, 4 and 5 (containing 976, 889 and 672 proteins, respectively) underwent significant increases in distinct waves: Proteins in Cluster 4 increased in levels during the MZT, reaching a peak at mid-embryogenesis and then declined somewhat thereafter. Proteins in Cluster 5 increased in levels after the MZT and continued to rise throughout the rest of embryogenesis. Proteins in Cluster 3 increased in levels at mid-embryogenesis and continued to increase thereafter. Strikingly, there was only one cluster that underwent a significant decrease in relative levels: 144 proteins were highest at the first time point and rapidly decreased to very low relative levels by the end of the MZT (Cluster 6).

Gene Ontology (GO) term analysis was conducted on each of the six clusters using the DAVID bioinformatics resource (da Huang et al., 2009a; da Huang et al., 2009b) (Supplemental file 2). Cluster 1, whose component proteins were at constant relative levels through the MZT and then decreased slightly during the remainder of embryogenesis, showed enrichment for terms related to the ubiquitin-proteasome system as well as core biological processes such as DNA replication and cell division. Cluster 2, whose proteins were at constant levels for the first part of the MZT, then increased slightly after ZGA and continued to gradually increase thereafter, was enriched for terms related to core components of transcription, splicing and translation. Enrichment of Clusters 1 and 2 for core cellular and molecular functions is consistent with the constant requirement for these components throughout development.

Cluster 4, which showed a rapid increase in relative protein levels during the MZT, was enriched for terms related to chromatin, sequence-specific DNA binding, transcriptional regulation, and cell fate specification. This is consistent with the fact that ZGA is known to be required to produce transcription factors that specify cell fate and pattern starting at the cellular blastoderm stage. Cluster 5, which increased in expression after the MZT, was enriched for terms related to cell adhesion, and several morphogenetic processes (tube formation, heart, neuromuscular junction), consistent with the sculpting of tissues and organs during this time window. Cluster 3, which increased during mid-to late-embryogenesis, was enriched for terms related to chitin production and secretion, myogenesis, and synaptogenesis, consistent with the final stages of embryonic development and the secretion of the larval cuticle prior to hatching from the egg.

Cluster 6, comprised of 144 proteins, was unusual in that relative expression was highest at the earliest stage of embryogenesis following which its proteins were rapidly degraded to low levels by the end of the MZT. These proteins must be either largely maternally supplied and already present in the oocyte, or rapidly synthesized from maternal mRNAs after egg activation or fertilization; we refer to them as ‘maternal’ proteins. This cluster showed enrichment for GO terms related to the eggshell as well as cytoplasmic ribonucleoprotein (RNP) granules, germ plasm and germ cell development (Supplemental file 2). Since these RNP components are present during the first phase of the MZT, when gene expression is regulated primarily post-transcriptionally, it is plausible that they participate in these processes. Furthermore, their degradation could serve to control the timing of these processes. The degradation of only a small fraction of maternally loaded proteins stands in striking contrast to maternally loaded mRNA species, two-thirds of which are degraded by the end of the MZT (Thomsen et al., 2010).

### Dynamics of the proteome relative to the transcriptome and translatome during the MZT

We next compared the relative changes in the proteome (this study) to those previously identified for the transcriptome and translatome (Eichhorn et al., 2016) at equivalent timepoints across the MZT. We focused on Cluster 4 (newly synthesized during the MZT) and Cluster 6 (degraded during the MZT).

A significant proportion of mRNAs encoding proteins in Cluster 4 underwent increases in RNA expression and translational efficiency at comparable timepoints (Fisher’s exact test *P* < 10^−16^ and *P* = 0.02, respectively; Figures 1D and F). However, whereas a significant proportion of the Cluster 6 unstable maternal proteins underwent a concomitant decrease in cognate mRNA levels, there was no association with changes in translation efficiency (Fisher’s exact test *P* < 10^−10^ and *P* = 0.26, respectively; Figures 1E and G). This last result is consistent with the fact that some of the Cluster 6 proteins, such as SMG, are synthesized during the MZT rather than during oogenesis; thus their cognate mRNAs exhibit high translational efficiency despite the fact that the protein is subsequently rapidly cleared (Eichhorn et al., 2016; Tadros et al., 2007). We note that a previous study that compared the embryonic proteome and transcriptome excluded almost all proteins in Cluster 6 from their analysis because of these proteins’ narrow expression window (Becker et al., 2018).

### Maternal ribonucleoprotein granule components are degraded during the MZT

A closer examination of the Cluster 6 RNP granule proteins revealed two temporally distinct classes. One subset was cleared from the embryo very rapidly and was depleted before the second timepoint (NC14); this subset includes the translational repressors Cup, TRAL and ME31B as well as the poly(A) polymerase, Wispy (Figure 1B). A second subset was degraded after NC14 and was depleted from the embryo by the third time point (germ band extension), at the end of the MZT. This subset includes SMG, the anterior determinant, Bicoid, germ plasm components such as Vasa, Oskar and Tudor (Figure 1C), as well as two subunits of the PNG kinase complex, PNG itself and a regulatory subunit, Plutonium (Supplemental file 1). We note that not all maternally loaded RBPs fall into Cluster 6; several were stable and fell into Clusters 1 or 2 (e.g., Brain tumor, Pumilio, Rasputin, Egalitarian, Belle, PABP, PABP2).

To verify the results of the mass spectrometry and to assess the dynamics of SMG, Cup, TRAL and ME31B expression at higher temporal resolution, we carried out Western blot analysis of extracts from embryos collected in overlapping 30 minute intervals through the first six hours of embryogenesis (Figure 2A, Figure 2 – Figure supplement 1). Cup, TRAL and ME31B are maternally supplied proteins with highest expression at the onset of embryogenesis (Sysoev et al., 2016; Wang et al., 2017). Our analysis showed that these proteins are rapidly degraded at 1–2 hours of embryogenesis, during the syncytial blastoderm stage, coincident with degradation of their cognate transcripts (Wang et al., 2017). Whereas Cup was completely cleared (0% remained at 4 hours), TRAL and ME31B decreased rapidly but then persisted at low levels (respectively, 6 and 13% remained). In contrast, SMG protein expression was low at the onset of embryogenesis and peaked at about 1 hour, corresponding with rapid translational derepression of the maternally loaded *smg* transcripts upon egg activation (Tadros et al., 2007). We note that this rapid increase was detectable in the high-resolution Western blot time-course but not in the lower resolution mass spectrometry time-course. SMG then underwent precipitous degradation at about 3 hours (Benoit et al., 2009), coinciding with blastoderm cellularization and high-level ZGA (4% of the maximum amount of SMG remained at 4 hours). Two additional proteins associated with the *nanos* repressor complex (Götze et al., 2017; Jeske et al., 2011) were also assessed: the DEAD-box helicase, Belle, and the cap-binding protein, eIF4E. Levels of Belle remained fairly constant (67% remained at 4 hours) whereas eIF4E gradually decreased in abundance but was clearly detectable throughout the time-course (19% remained after 4 hours). These Western blot analyses were fully consistent with the proteomic data presented above as well as previous reports (Sysoev et al., 2016; Wang et al., 2017).

**Figure 2.**
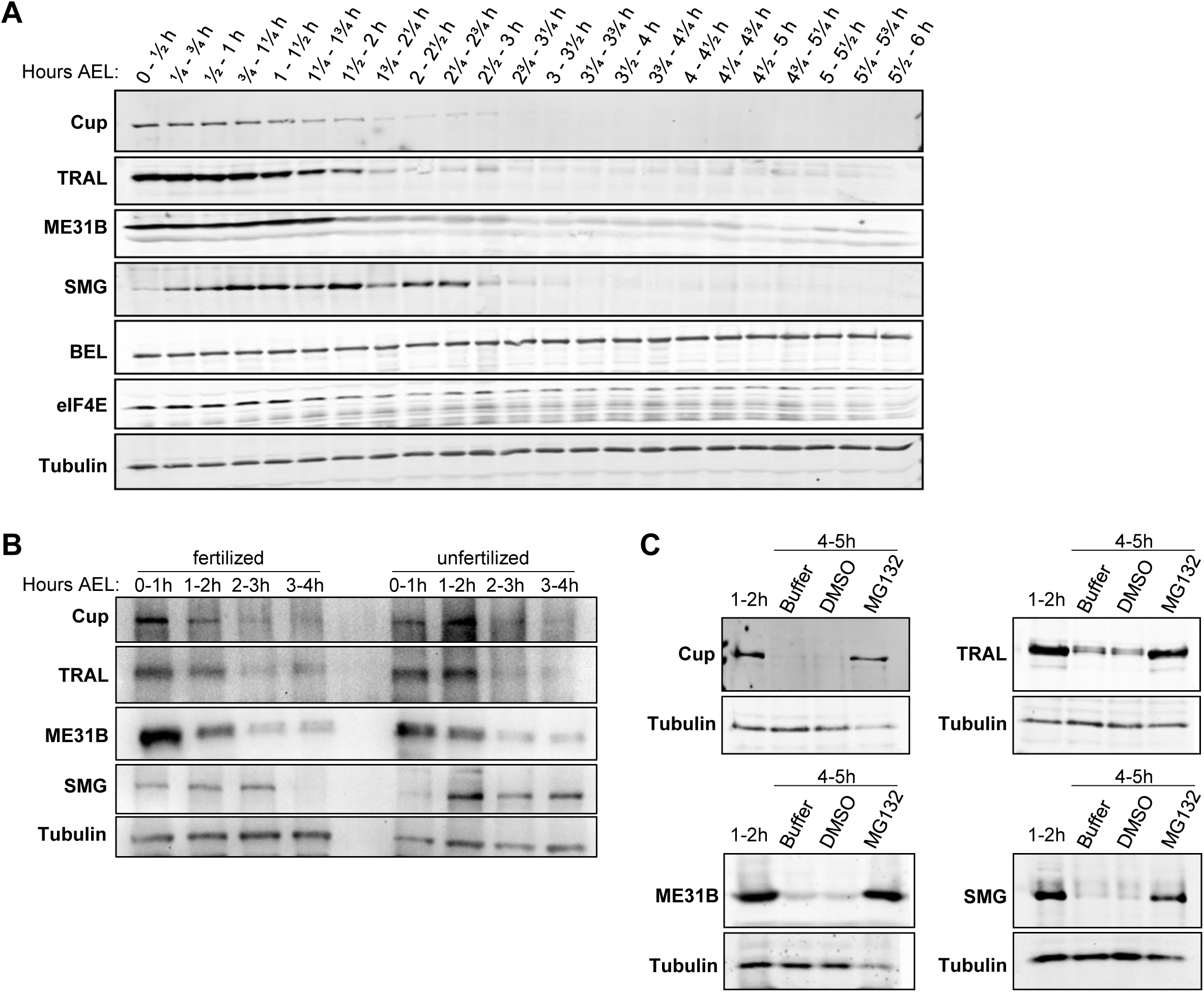
SMG, Cup, TRAL and ME31B are degraded at distinct times during the MZT through the ubiquitin proteasome system. **A.** Developmental western blot of wild-type embryos collected in 30-minute time windows and aged at 15-minute intervals over the first six hours after egg-lay (AEL). Cup, TRAL and ME31B are depleted by around 1.5 hours AEL, while SMG levels increase in the early embryo and are subsequently cleared by about 2.5 hours AEL. BEL and eIF4E remain expressed for the duration of the time course. Tubulin was probed as a loading control. See **Figure 2 – figure supplement 1** for quantification. **B.** Developmental western blot of wild-type embryos collected over the first 4 hours of embryogenesis from mated females (fertilized) and unmated females (unfertilized). Degradation of Cup, TRAL and ME31B are unaffected in unfertilized eggs, while SMG protein fails to be degraded by 3-4h AEL. **C.** Western blots of 1-2h old embryos that were permeabilized and incubated for three hours in buffer, DMSO control, or 100μM MG132 and aged to 4-5h AEL. All four RBPs shown are stabilized by MG132 treatment.

Previous studies have shown that the clearance of SMG protein from the embryo depends on ZGA (Benoit et al., 2009). To determine the role of ZGA in the degradation of all four repressors, we analyzed the expression of SMG, Cup, TRAL, and ME31B in activated, unfertilized eggs, which carry out maternally directed post-transcriptional and post-translational processes, but do not undergo transcriptional activation (Bashirullah et al., 1999; Page and Orr-Weaver, 1997; Tadros et al., 2003). Embryo extracts were collected from wild-type mated and unmated females in one-hour intervals over four hours, and expression of the RBPs was assayed by Western blotting (Figure 2B). The clearance of Cup, TRAL and ME31B was unaffected in unfertilized eggs, indicating that their degradation is not dependent on ZGA. In contrast, SMG protein was stable in unfertilized eggs, confirming that degradation of SMG requires zygotically expressed factors (Benoit et al., 2009). Thus, Cup-TRAL-ME31B and SMG differ both in the timing of their degradation and in whether they require zygotically synthesized gene products to accomplish this process.

### Degradation of the repressors is regulated by the ubiquitin-proteasome system

To determine whether clearance of SMG, Cup, TRAL and ME31Bs occurs via the ubiquitin-proteasome system, we inhibited the proteasome in developing embryos using the small molecule, MG132. 1–2 hour-old embryos were permeabilized (Rand et al., 2010) and incubated with either MG132 or control buffer, then allowed to develop for a further three hours. Western blots on controls showed that SMG, Cup, TRAL, and ME31B were degraded as expected. Strikingly, treatment with 100μM MG132 resulted in stabilization of all four RBPs (Figure 2C).

If the ubiquitin-proteasome system indeed targets these proteins for degradation, it should be possible to identify specific sites of ubiquitination. In MS analysis of tryptic protein digests, ubiquitination is visible as a lysine to which the two C-terminal glycine residues of ubiquitin are attached; often, the modified lysine residue is not cleaved by trypsin. An analysis of our previous MS data (Götze et al., 2017) revealed ubiquitination signatures in all four repressors (Supplemental file 3). To identify additional ubiquitination sites on these proteins, we performed immunoprecipitation coupled to mass spectrometry (IP-MS) on embryo extract prepared in the presence of three inhibitors: the proteasome inhibitor, MG132; the inhibitor of deubiquitinating enzymes, PR-619; and EDTA to block all ATP-dependent processes. These conditions are expected to stabilize ubiquitinated proteins. Anti-Cup antibody was used to IP the repressor complex, and additional ubiquitination sites were identified by MS (Supplemental file 3).

Together these data strongly support a role for the ubiquitin-proteasome in clearance of SMG, Cup, TRAL and ME31B proteins during the MZT.

### Two distinct ubiquitin-ligase complexes interact with the repressors

To identify the specific E3 ligases that regulate degradation of the repressors during the MZT, we performed IP-MS from transgenic embryos expressing FLAG-tagged SMG protein. As expected, Cup, TRAL and ME31B, as well as BEL and eIF-4E were highly enriched in FLAG-SMG IPs relative to control IPs, placing them among the top interactors (Figure 3A, Supplemental file 4), recapitulating the known repressive complex (Götze et al., 2017; Jeske et al., 2011). In addition, we captured RNA-independent interactions with several members of two distinct multi-subunit E3 ubiquitin ligase complexes: the SCF complex and the CTLH complex. SCF complex components that were identified included core components such as SKPA (Skp) and CUL1 (Cullin), as well as SLMB and CG14317 (two F-box proteins) (Ravid and Hochstrasser, 2008) while the CTLH complex was represented by all characterized *Drosophila* subunits (Francis et al., 2013), including RanBPM, Muskelin, CG6617, CG3295, CG7611 and CG31357 (Figure 3B, Supplemental file 4). Detection of the CTLH complex confirmed earlier data (Götze et al., 2017).

**Figure 3.**
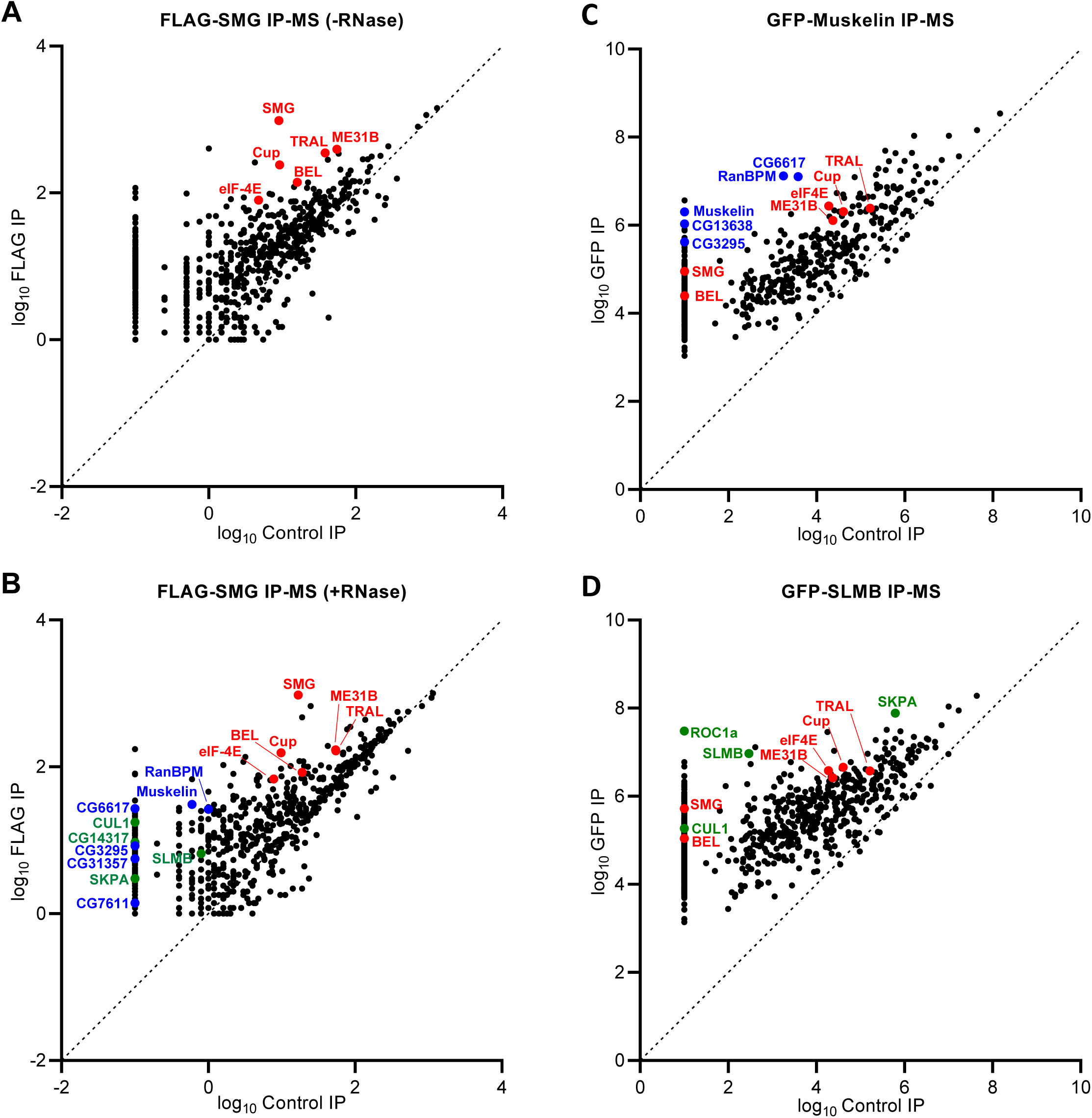
SMG interacts with repressor RBPs and two distinct E3 ubiquitin ligase complexes. **A-B.** FLAG IP-MS of 0-3h embryo lysate collected from transgenic flies expressing FLAG-SMG, and homozygous for the deletion allele *smg*^*47*^. FLAG IP from non-transgenic embryo lysate was used as control. Average spectral counts are plotted for proteins detected at ≥1 in FLAG-SMG IP on average across at least 4 biological replicates. **A.** In the absence of RNase A, SMG interacts strongly with its co-repressors (red). **B.** IP in the presence of RNase A captured RNA-independent interactions with two E3 ubiquitin ligase complexes: the CTLH complex (blue: Muskelin, RanBPM, CG6617, CG3295, CG31357 and CG7611), and the SCF complex (green: CUL1, SKPA, CG14317 and SLMB). **C-D.** GFP IP-MS of 0-2h embryo lysate from embryos expressing either GFP-Muskelin or GFP-SLMB. GFP IP from non-transgenic embryo lysate was used as control. Average iBAQ intensities (Cox and Mann, 2008) for proteins detected across 3 biological replicates are plotted. **C.** GFP-Muskelin interacts with the RBPs and other members of the CTLH complex. **D.** GFP-SLMB interacts with the RBPs and other members of the SCF complex. Note: Cutoffs were not applied to negative controls. Proteins not detected in control IPs are assigned an average spectral count of 0.1 (A-B), or an iBAQ value of 10 (C-D) to avoid log(0); these small values are at least twofold less than the lowest detected values across all experiments.

To verify the interaction of these two E3 ligase complexes with the repressors, we performed reciprocal IP-MS experiments using embryos expressing GFP-SLMB or GFP-Muskelin. GFP pull-downs from RNase-treated extracts captured enrichment for additional subunits of the respective complexes, as well as all four repressors relative to control (Figure 3C-D, Supplemental file 5). The association of all four proteins with both E3 ligase complexes presumably reflects their stable complex formation.

### Degradation of Cup, TRAL and ME31B is directed by the CTLH E3 ubiquitin ligase

To investigate the role of the SCF and the CTLH E3 ubiquitin ligase complexes in degradation of the repressors during the MZT, we performed maternal RNAi knockdown experiments for several core members of the SCF and CTLH complexes. Embryos were collected at one-hour intervals from female flies with germline knockdown, and expression of SMG, Cup, TRAL and ME31B was quantified by Western blot over the first four hours of embryogenesis. Knockdown was confirmed in these embryos by RT-qPCR and, in the case of SLMB, also by western blot (Figure 4 – Figure supplement 1; Figure 5 – Figure supplement 1).

**Figure 4.**
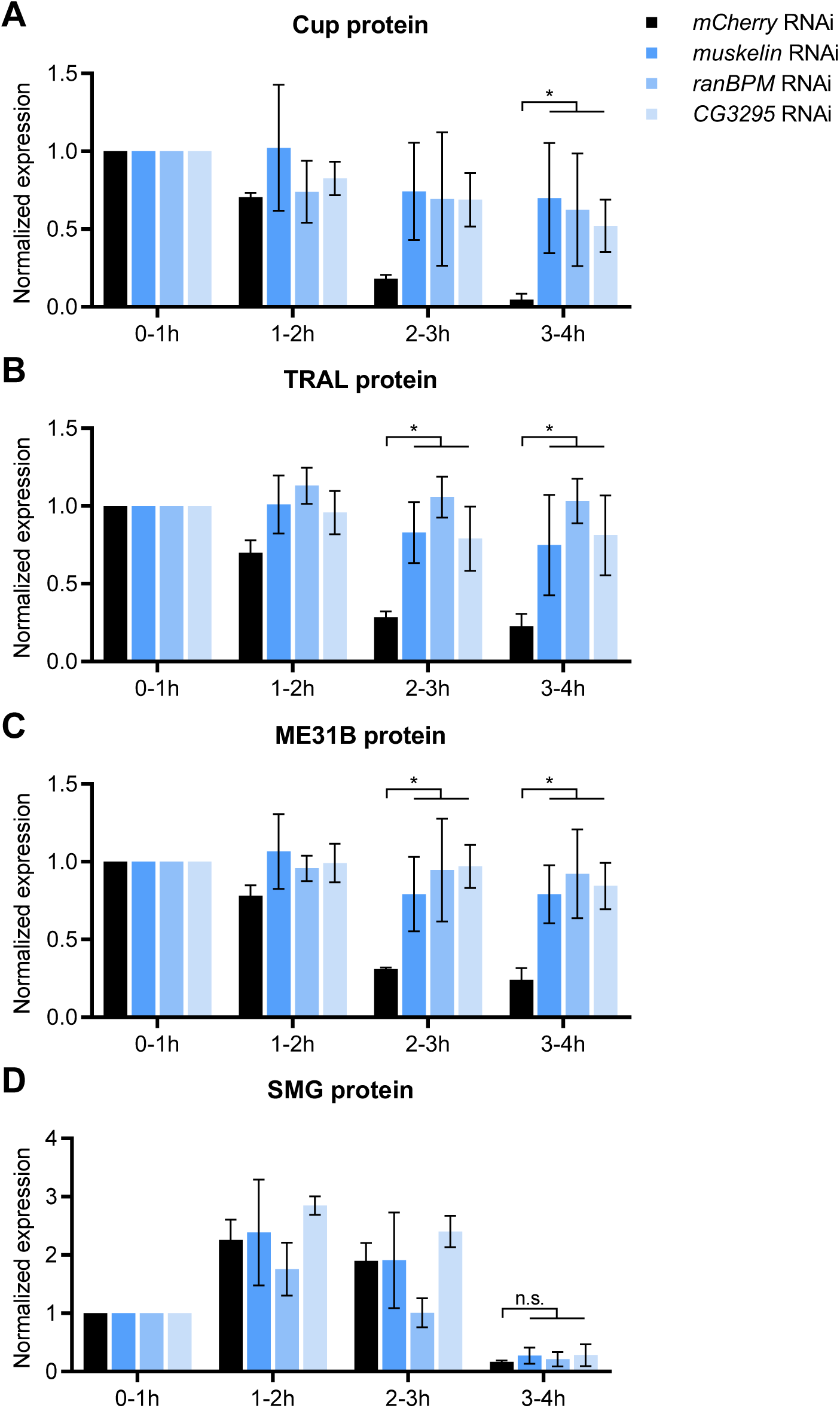
The CTLH complex directs the degradation of Cup, TRAL, and ME31B but not SMG. Quantified developmental western blots of RBP expression. Embryos were collected from maternal knockdown of CTLH complex members over the first four hours after egg-lay. Knockdown of *muskelin, ranBPM* and *CG3295* each independently resulted in significant stabilization of Cup (**A**), TRAL (**B**) and ME31B (**C**) relative to control *mCherry* knockdown. **D.** SMG protein degradation was unaffected by knockdown of the CTLH complex. \**P* < *0.05, n.s. =* not significant, n = 3, error bars = SD, *S*tudent’s *t*-test. Knockdown was confirmed by qPCR (**Figure 4 – figure supplement 1**), see also **Figure 4 – figure supplement 2-3**.

**Figure 5.**
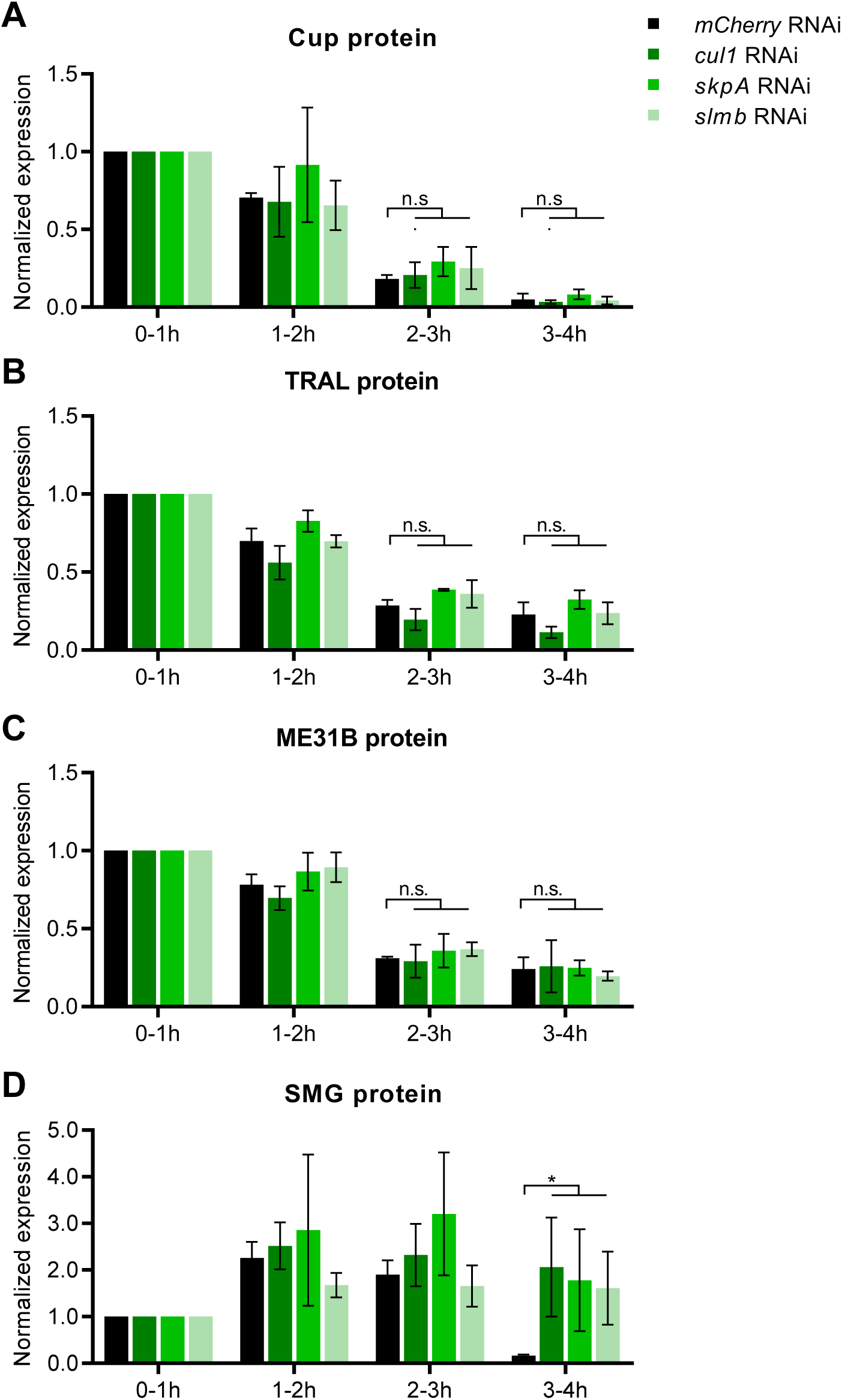
The SCF complex directs the degradation of SMG but not Cup, TRAL and ME31B. **A-D.** Quantified developmental western blots of RBP expression. Embryos were collected from maternal knockdown of SCF complex members over the first four hours after egg-lay. Cup (**A**), TRAL (**B**) and ME31B (**C**) protein degradation was unaffected by knockdown of the SCF complex. **D.** Knockdown of *cul1, skpA* and *slmb* each independently resulted in significant stabilization of SMG protein relative to control *mCherry* knockdown \**P* < *0.05*, n.s. *=* not significant, n = 3, error bars = SD, *S*tudent’s *t*-test. Knockdown was confirmed by qPCR (**Figure 5 – figure supplement 1**), see also **Figure 5 – figure supplement 2**.

Maternal knockdown of the CTLH complex members Muskelin, RanBPM, and CG3295 resulted in significant stabilization of Cup, TRAL and ME31B proteins (Figure 4A-C), while SMG protein degradation was completely unaffected (Figure 4D). In these experiments, knockdown of the CTLH complex components also resulted in a delay in the degradation of *cup, tral* and *me31B* transcripts (Figure 4 – Figure supplement 2). DAPI staining of the RNAi embryos revealed a developmental delay in a subset of embryos (about 20%). To eliminate the possibility that developmentally delayed or arrested embryos contributed to repressor protein ‘persistence’, we visualized embryo development live under halocarbon oil and picked embryos at three specific developmental stages during the MZT for analysis: Stage 2 (about 0.5–1 hr AEL), Stage 5b (about 2.5 hr AEL), and Stage 7b (about 3 hr AEL). Developmentally delayed or abnormal embryos were thus excluded by this method. Knockdown of any of the three members of the CTLH complex led to nearly complete stabilization of Cup, TRAL and ME31B through to Stage 7b whereas, in controls, all three proteins were degraded by Stage 5b (Figure 4 – Figure supplement 3A-D). RT-qPCR quantification of mRNA expression in these staged embryos showed that *cup* mRNA was cleared from the embryo by Stage 7b both in knock-down and control embryos. This result excludes new synthesis as the reason for the persistence of Cup under conditions of CTLH complex knock-down, thus providing strong evidence that the CTLH complex is required for degradation of the Cup protein (Figure 4 – Figure supplement 3E). However, *tral* and *me31B* transcripts were partially stabilized in Stage 7b embryos (Figure 4 – Figure supplement 3F,G); thus we cannot rule out the possibility that translation from their stabilized cognate transcripts contributes in part to the persistence of TRAL and ME31B protein. Either way, it is clear that the CTLH plays a key role in clearance of Cup and, likely, also TRAL and ME31B, but not in clearance of SMG.

### Degradation of SMG is directed by the SCF E3 ubiquitin ligase

In direct contrast to Cup, TRAL and ME31B, we found that maternal knockdown of the SCF complex members CUL1, SKPA or SLMB had no effect on Cup, TRAL and ME31B expression but resulted in reproducible stabilization of SMG protein (Figure 5). Furthermore, we confirmed by RT-qPCR that degradation of *smg* mRNA, which is coincident with SMG protein’s clearance, was not affected in these embryos (Figure 5 – Figure supplement 2A). Thus, persistence of SMG in the absence of the SCF E3-ligase complex must have been a result of protein stabilization. Finally, to exclude the possibility that stabilization of SMG was a result of a developmental delay resulting from these knockdowns, we performed immunostaining of SMG in the RNAi embryos. Whereas SMG normally disappears from the bulk cytoplasm of embryos prior to gastrulation, in knockdown of core members of the SCF complex, we found gastrulating embryos ubiquitously staining for high levels of SMG protein (Figure 5 - Figure Supplement 2B).

Taken together, these data provide strong evidence that the SCF E3 ubiquitin ligase complex regulates degradation of SMG protein at the end of the MZT whereas the CTLH complex directs degradation of Cup, TRAL and ME31B.

### Temporal regulation of the E3 ligase complexes

What determines the timing of E3 ligase complex action? We assessed the dynamics of CTLH and SCF subunits in our TMTc+ MS data and found that, with the exceptions described below, the subunits detected (CTLH: RanBPM, CG3295, CG6617, CG7611, CG31357; SCF: CUL1, SKPA, ROC1a, SLMB) were present throughout embryogenesis (Figure 6). We note that these results are consistent with Cluster 1 proteins, which are present at almost constant relative levels throughout embryogenesis, showing enrichment for the GO term ‘ubiquitin-proteasome system’.

**Figure 6:**
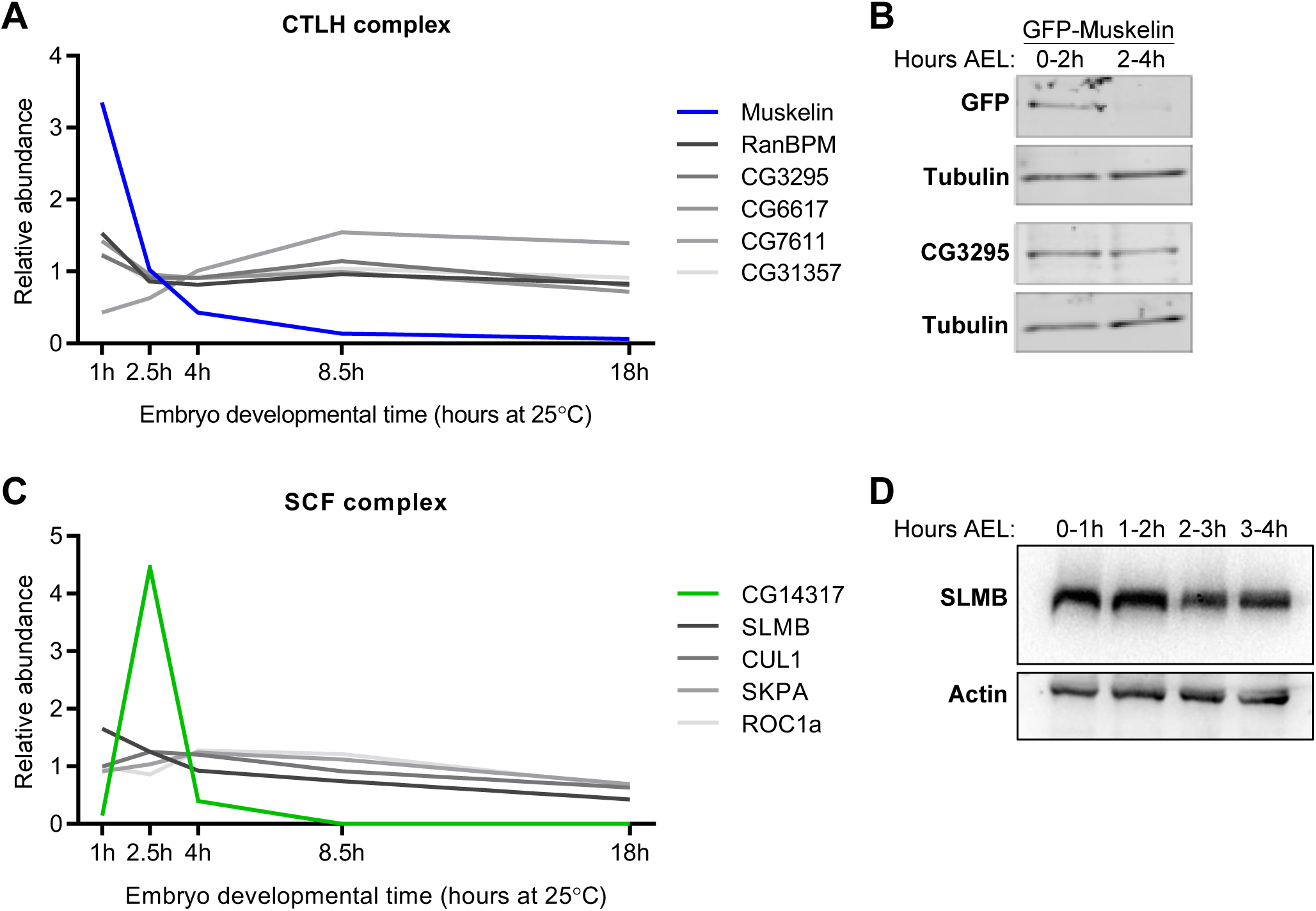
Subunits of the E3 ligase complexes are temporally regulated during the MZT. **A.** Expression of subunits of the CTLH complex captured in the developmental proteome. Most subunits have relatively constant levels throughout embryogenesis, while levels of Muskelin are highest at the first time point and then decrease rapidly. **B.** Western blot of embryos expressing GFP-Muskelin. Anti-GFP (top) confirmed rapid clearance of GFP-Muskelin from the embryo. Anti-CG3295 (bottom) confirmed its stable expression during the MZT. **C.** Expression of subunits of the SCF complex captured in the developmental proteome. Most subunits exhibit relatively constant levels throughout embryogenesis, while levels of the F-box subunit, CG14317 increased rapidly during the MZT and then decreased very rapidly by the end of the MZT, with peak levels coinciding with degradation of SMG protein. **D.** Developmental western blot of control RNAi embryos, confirming the stable expression of SLMB during the MZT. Note: The same blot shown here is used to confirm SLMB knockdown in Figure 5 – figure supplement 1.

In contrast to the other CTLH subunits, the relative level of Muskelin decreased rapidly during the MZT (Figure 6A), which we confirmed by Western blotting of GFP-tagged Muskelin and comparison to endogenous CG3295 (Figure 6B). Furthermore our Cup IP-MS to map ubiquitination sites on the repressors revealed that Muskelin is ubiquitinated in early embryos (Supplemental file 3). Thus, the dynamics of Muskelin clearance would restrict CTLH function to the early phase of the MZT, fully consistent with the timing of Cup, TRAL and ME31B degradation.

For the SCF complex, our TMTc+ MS showed that the CUL1, SKPA, ROC1a and SLMB subunits were present at relatively constant levels throughout embryogenesis (Figure 6C) and Western blots confirmed that SLMB is expressed throughout the MZT (Figure 6D). The F-box subunit confers substrate-specificity to the SCF complex, and SLMB is one of two F-box proteins that were found to interact with SMG by IP-MS (Figure 3B). Strikingly, the second F-box protein, CG14317, displayed a highly unusual expression pattern: the CG14317 mRNA is absent at the beginning of embryogenesis, is zygotically expressed, peaking during the late MZT, and then drops precipitously (Brown et al., 2014; Graveley et al., 2011). The mRNA is not detected at any other stage of *Drosophila* development, neither is it found in any tissue or cell line that has been profiled (http://flybase.org/reports/FBgn0038566). Our MS data showed that the CG14317 protein follows an almost identical expression pattern to that of its cognate mRNA: absent at the first timepoint, peak expression at the second timepoint (NC14), sharp drop at the third timepoint (Figure 6C). We have not yet assessed CG14317 function by RNAi; however, the extremely narrow time-window of CG14317 expression coincides almost exactly with when SMG protein degradation is triggered. Furthermore, degradation of SMG near the end of the MZT depends on zygotic transcription and CG14317 mRNA is produced zygotically. These data lead us to speculate that CG14317 could serve as a precise timer for SCF action on SMG (see Discussion). The fact that knockdown of SLMB stabilizes SMG protein suggests that both F-box proteins are necessary for SMG degradation, with CG14317 serving as the timer.

### Persistent SMG protein downregulates zygotic re-expression of its target transcripts

Finally, we turned to possible biological functions of repressor clearance with a focus on SMG. The SMG RBP contains a dimerization domain near its N-terminus (Tang et al., 2007) and a SAM-PHAT RNA-binding domain towards the C-terminus (Aviv et al., 2006; Aviv et al., 2003). We produced transgenic fly lines expressing FLAG-P53-tagged SMG, either full-length (‘FLAG-SMG’) or missing the C-terminal 233 amino acids (‘FLAG-SMG767Δ999’) (Figure 7A-B). The transgenes were under control of the endogenous *smg* gene’s regulatory elements and, to avoid potential complications resulting from dimerization of transgenic FLAG-SMG with endogenous SMG, all transgenic protein expression was assayed in the *smg*^*47*^ null-mutant background (Chen et al., 2014). We found that truncation of the C-terminal 233 amino acids led to stabilization of the protein throughout the MZT (Figure 7B). This positioned us to assess the effects of persistent SMG.

**Figure 7:**
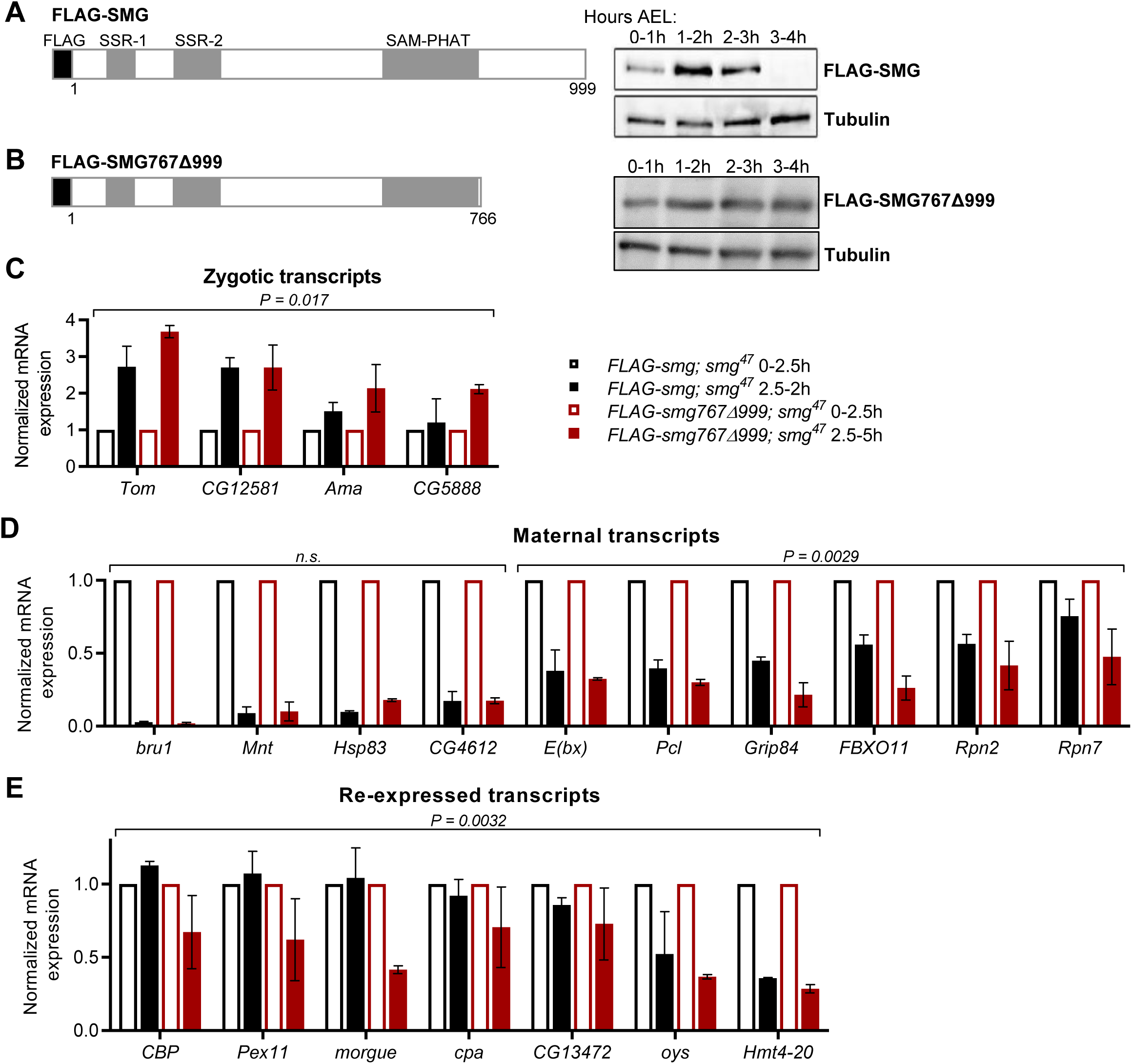
Persistent SMG protein downregulates zygotic re-expression of its target transcripts. **A-B.** Transgenic flies were generated expressing either FLAG-tagged full-length SMG or SMG767Δ999 truncated C-terminal to its SAM-PHAT RNA-binding domain. Transgenes were under the control of endogenous regulatory elements. Developmental western blots were performed on embryos collected from transgenic flies in the *smg*^*47*^ deletion-mutant background **A**. FLAG-SMG expression resembled that of endogenous SMG. **B**. SMG767Δ999 protein was stabilized and remained through the MZT. **C-E.** Embryos were collected from the transgenic flies at two time points during the MZT, and gene expression was assayed by RT-qPCR. **C**. Expression of transcripts that depend on SMG for zygotic transcription was rescued by SMG767Δ999 to similar or higher levels than by full-length SMG. These transcripts are predicted not to be direct targets for SMG-binding since they have low SRE scores (see Methods). **D.** Degradation of SMG-bound target maternal transcripts was rescued by SMG767Δ999. Where transcripts were not completely degraded by FLAG-SMG, SMG767Δ999 persistence resulted in further degradation of these targets to significantly lower levels (rightmost 6 genes). **E.** SMG-target maternal transcripts that are re-expressed zygotically were significantly downregulated in their zygotic levels by persistence of SMG767Δ999. Wilcoxon signed-rank test *P*-values shown for each gene group; two biological replicates for each gene, error bars = SD.

SMG is known to target hundreds of maternal transcripts for degradation during the MZT (Chen et al., 2014; Tadros et al., 2007). A subset of these transcripts is re-expressed upon ZGA (Benoit et al., 2009; De Renzis et al., 2007; Thomsen et al., 2010) and the CCR4-NOT deadenylase complex remains expressed at fairly constant levels throughout embryogenesis (Cluster 1, and see Temme et al., 2004). Thus, we hypothesized that persistent SMG protein may result in unintentional targeting of zygotically re-expressed SMG targets. To test this hypothesis we compared transcript dynamics in *smg*^*47*^ mutant embryos expressing either full-length SMG or SMG767Δ999. Gene expression was assayed by RT-qPCR in 0–2.5 hour embryos (representing the maternal phase of the MZT), and 2.5–5 hour embryos (representing the zygotic phase of the MZT).

Embryos that lack SMG protein fail to undergo both maternal mRNA decay and ZGA (Benoit et al., 2009; Tadros et al., 2007). We, therefore, first carried out two control experiments to assess whether SMG767Δ999 rescues these processes. One control was strictly zygotic transcripts that lack SMG-binding sites (Benoit et al., 2009). We found that these were expressed at comparable (or even higher; Wilcoxon signed rank test *P* < 0.02) levels for SMG767Δ999 versus full-length SMG, showing that SMG767Δ999 rescues ZGA (Figure 7C). A second control was degradation of SMG-bound maternal transcripts that are targeted by SMG but are not re-expressed zygotically (Chen et al., 2014; Tadros et al., 2007). We found that these were degraded in SMG767Δ999 embryos, sometimes (*Frip84, FBXO11, Rpn2, Rpn7*) to even lower levels than in embryos expressing full-length SMG (Wilcoxon signed rank test *P* < 3×10^−3^; Figure 7D). Thus SMG767Δ999 also rescues maternal mRNA decay. Finally we assayed the expression of transcripts that met all of three criteria: (1) maternally supplied and degraded in a SMG-dependent manner (Tadros et al., 2007); (2) directly bound by SMG (Chen et al., 2014) ; and (3) zygotically re-expressed at high levels shortly after ZGA (Benoit et al., 2009). Strikingly, *smg*^*47*^ embryos expressing SMG767Δ999 showed significantly lower zygotic expression of these transcripts when compared to *smg*^*47*^ embryos expressing full-length SMG (Wilcoxon signed rank test *P* < 4×10^−3^; Figure 7E).

We conclude that precise temporal regulation of SMG protein degradation at the end of the MZT is required to permit proper zygotic re-expression of transcripts with SMG-binding sites.

## DISCUSSION

During the MZT, a massive degradation of the maternal mRNA transcriptome occurs. Here we have shown that, in contrast to the maternal transcriptome, an extremely small subset of the maternal proteome is cleared. Furthermore, the cleared proteins are enriched for RNP granule components. This is consistent with the importance of post-transcriptional processes during the first, ‘maternal’, phase of the MZT, and the possible need to downregulate these processes upon ZGA and the switch to zygotic control of development. By focusing on a subset of these RNP components, which function as post-transcriptional repressors, we have uncovered precise temporal control of their clearance by two distinct E3 ubiquitin ligase complexes. Intriguingly, SMG is degraded at a later time during the MZT than its co-repressors Cup, TRAL and ME31B. The SCF E3 ligase governs the degradation of SMG, whereas the CTLH E3 ligase is responsible for the degradation of Cup, TRAL and ME31B. We have also shown that clearance of SMG is essential for appropriate levels of re-expression of a subset of its targets during ZGA. Our results raise questions about how temporal specificity of protein degradation is regulated, as well as why at least two temporally distinct mechanisms of protein degradation exist during the MZT.

Previous studies have suggested that the maternal proteome may behave very differently from the maternal transcriptome during the MZT. For example, in *C. elegans*, a quarter of the transcriptome is downregulated whereas only 5% of the proteome shows a similar decrease (Stoeckius et al., 2014). In frog embryos there is also a discordance between the temporal patterns of protein and mRNA, although protein dynamics can be reasonably well predicted when taking into consideration the absolute concentration of the maternal protein load in the unfertilized egg, and modeling protein synthesis and degradation (Peshkin et al., 2015).

Among the best studied of the RNP granule components that we have shown to be rapidly cleared during the MZT are several post-transcriptional repressors of translation and mRNA stability. Here we have presented evidence that two E3 ligase complexes, CTLH and SCF, target different repressors at different times during the *Drosophila* MZT: respectively Cup-TRAL-ME31B early in the MZT and SMG at the end of the MZT. We have also presented protein expression data that support the hypothesis that timing of E3 ligase function is at least in part determined by the timing of expression of one or more of their component subunits, notably Muskelin for CTLH and CG14317 for SCF.

Regarding Muskelin, we have shown that during the *Drosophila* MZT, while most components of the CTLH complex display constant expression levels, Muskelin protein is degraded with a similar profile to its target repressors. Mammalian Muskelin has been shown to be auto-ubiquitinated and targeted for degradation (Maitland et al., 2019). Our detection of a ubiquitinated peptide on Muskelin supports the possibility that the *Drosophila* CTLH complex may also be negatively auto-regulated through its Muskelin subunit during the *Drosophila* MZT. We note that, going from Stage 14 oocytes to activated eggs or early (0–1 hour) embryos, previous studies have shown that there are no significant changes in either the levels of CTLH subunit proteins (including Muskelin) or the translation index of their cognate transcripts (Eichhorn et al., 2016; Kronja et al., 2014b). Thus we speculate that post-translational modification of one or more CTLH subunits must activate CTLH function at the beginning of the MZT. The degradation of Cup, TRAL and ME31B depend on the PNG kinase (Wang et al., 2017), which itself has temporally restricted activity coinciding with degradation of these repressors (Hara et al., 2017). PNG-dependent phosphorylation of Cup, TRAL and ME31B may make them ubiquitination substrates. Concomitant PNG-dependent activation of the CTLH complex, coupled with subsequent self-inactivation of the complex through Muskelin degradation, would provide a precise time window for CTLH function and, therefore, for degradation of Cup, TRAL and ME31B early in the MZT.

In contrast to these three co-repressors, degradation of SMG occurs near the end of the MZT and depends on zygotic gene expression. While the levels of most SCF complex subunits are constant during the MZT, the F-box protein, CG14317, displays a unique expression pattern: CG14317 protein and mRNA are absent at the beginning of the MZT, are zygotically synthesized, peak in NC14 embryos, and sharply decline shortly thereafter. This extremely narrow time window of CG14317 expression coincides almost perfectly with the timing of SMG protein degradation and, coupled with the zygotic nature of its accumulation, makes CG14317 a strong candidate to be a timer for SCF function. At present there are no forward- or reverse-genetic reagents available to test this hypothesis. The fact that knockdown of SLMB stabilizes SMG protein suggests that both F-box proteins are necessary for SMG degradation, with CG14317 serving as the timer.

Since Cup, TRAL and ME31B are known to function as co-repressors in a complex with SMG (Götze et al., 2017; Jeske et al., 2011), why is the timing of degradation of Cup-TRAL-ME31B and SMG differentially regulated? While the SMG-Cup-TRAL-ME31B-mRNA complex has been characterized to be extremely stable *in vitro* (Jeske et al., 2011), it would be disrupted *in vivo* by the degradation of Cup, TRAL and ME31B (or by the degradation of *nos* and other target mRNAs). SMG directs translational repression both through AGO 1 and through Cup, TRAL and ME31B, as well as transcript degradation through recruitment of the CCR4-NOT deadenylase (Chen et al., 2014; Nelson et al., 2004; Pinder and Smibert, 2013; Semotok et al., 2005; Tadros et al., 2007; Zaessinger et al., 2006). CTLH-driven degradation of Cup, TRAL and ME31B would abrogate SMG-Cup-TRAL-ME31B-dependent translational repression but not AGO1-dependent repression, since AGO1 levels increase during the MZT (Luo et al., 2016). The relative contributions of AGO1 versus Cup-TRAL-ME31B to translational repression by SMG are unknown. That said, the CCR4-NOT deadenylase is present both during and after the MZT (Temme et al., 2004); thus, SMG-dependent transcript degradation would occur both before and after clearance of Cup, TRAL and ME31B. 12% of SMG-associated transcripts are degraded but not repressed by SMG (Chen et al., 2014). Perhaps this subset is bound and degraded by SMG late in the MZT, after the drop in Cup, TRAL and ME31B levels.

Another possible role for clearance of ME31B and TRAL derives from studies in budding yeast, where it has been shown that their orthologs, respectively Dhh1p and Scd6p, have a potent inhibitory effect on ‘general’ translation (Coller and Parker, 2005; Nissan et al., 2010; Rajyaguru et al., 2012). If this is also true in *Drosophila*, then degradation of ME31B and TRAL, which are present at exceedingly high concentrations in embryos (Götze et al., 2017), might also serve to permit high-level translation during the second phase of the MZT, as previously hypothesized (Wang et al., 2017).

We previously showed that SMG has both direct and indirect roles in the MZT. SMG’s direct role is to bind to a large number of maternal mRNA species and target them for repression and/or degradation (Chen et al., 2014; Tadros et al., 2007). Two indirect effects become apparent in *smg* mutants: First, if maternal transcripts fail to be degraded and/or repressed, ZGA fails or is significantly delayed, likely because mRNAs encoding transcriptional repressors persist (Benoit et al., 2009; Luo et al., 2016). Second, since zygotically synthesized microRNAs direct a second wave of maternal mRNA decay during the late-MZT, in *smg* mutants failure to produce those microRNAs results in failure to eliminate a second set of maternal transcripts late in the MZT (Benoit et al., 2009; Luo et al., 2016).

Here we have uncovered a role for rapid clearance of the SMG protein itself late in the MZT: to permit zygotic re-expression of a subset of it targets. Notably, stabilized SMG (SMG767Δ999) rescues both clearance of its maternal targets and ZGA, excluding the possibility that lower that normal levels of re-expressed targets is a result of defective SMG function upon deletion of its C-terminus. Indeed, in our control experiments, SMG’s exclusively maternal targets actually drop to lower levels than normal, likely because SMG767Δ999 continues to direct their decay beyond when SMG normally disappears from embryos. Furthermore, in our other control, strictly zygotic transcripts that lack SMG-binding sites are expressed at higher levels in SMG767Δ999-rescued mutants than in full-length-SMG-rescued mutants. This result is consistent with the hypothesis that clearance of transcriptional repressors by SMG permits ZGA (Benoit et al., 2009); persistent SMG would clear these repressors to lower levels than normal, hence resulting in higher zygotic expression. The higher-than-normal expression of zygotic transcripts that lack SMG-binding sites makes the lower-than-normal levels of SMG’s zygotically re-expressed target transcripts by SMG767Δ999 even more striking. Together these data support a model in which the timing of both SMG synthesis and clearance is important for orderly progression of the MZT.

## MATERIALS AND METHODS

### Fly Strains

Flies were cultivated under standard laboratory conditions at 25°C. Wild-type strains included Canton S, *w*^*1118*^ and *y w*. Strains for GFP IP-MS experiments were: *UAS:GFP-Slimb-6 / CyO* (gift from Daniel St Johnston, Cambridge); *w*; P{UAS-muskelin.GFP}attP2* (Bloomington Drosophila Stock Center [BDSC] #65860); *nos-GAL4: w*^*1118*^; *P{GAL4::VP16-nos.UTR}CG6325*^*MVD1*^ (gift from Martine Simonelig, Montpellier, BDSC #4937). RNAi strains against proteins of interest were from the Transgenic RNAi Project (TRiP) and obtained from BDSC: Muskelin (#51405), RanBPM (#61172), CG3295 (#61896), CUL1 (#36601), SKPA (#32991), SLMB (#33898). mCherry RNAi was used as control (gift from T. Hurd, BDSC #35785). The maternal-GAL4 driver used was *y*^*1*^ *w*; P{matalpha4-GAL-VP16}67; P{matalpha4-GAL-VP16}1*5 (BDSC #80361).

### Primary antibodies

Primary antibodies used were as follows: Rat anti-Cup (Nakamura et al., 2004), rabbit anti-ME31B and rabbit anti-TRAL (Nakamura et al., 2001) were gifts from A. Nakamura. A second rabbit anti-ME31B (Harnisch et al., 2016) and rat anti-TRAL (Götze et al., 2017) were also used. Additional antibodies included guinea pig anti-SMG (Tadros et al., 2007), rabbit anti-SMG (Chartier et al., 2015), and rabbit anti-BEL (Götze et al., 2017). Guinea pig anti-SLMB was a gift from G.C. Rogers (Brownlee et al., 2011). For Cup-IP, a rabbit antibody directed against amino acids 1103-1117 of Cup (Eurogentec) was used. Anti-eIF4E and anti-CG3295 antibodies were raised in rat (Eurogentec). The protein eIF4E1, isoformA, was expressed in *E. coli* as a His6-SUMO fusion protein. After metal affinity chromatography, the N-terminal SUMO domain was cleaved off with ULP protease, and the two protein fragments were separated by a second metal affinity column. CG3295 was expressed in *E. coli* as a His6 fusion protein. The protein was purified from inclusion bodies under denaturing conditions via metal affinity chromatography. Commercially available mouse anti-Tubulin (Sigma) and mouse anti-Actin (Sigma) were also used as western loading controls.

### Embryo collection

Embryos were collected and aged at 25°C unless otherwise indicated. Cages containing male and female adult *Drosophila* were set up with apple juice agar plates supplemented with yeast paste. For unfertilized egg collection, unmated females were used, and males were housed in an adjacent cage separated by mesh to promote egg-lay. After egg-lay and development to the desired age, excess yeast was scraped off, and embryos were dechorionated with cold 4.2% sodium hypochlorite for 1–2 minutes, collected on a nylon mesh, and rinsed with PBST (PBS, 0.1% Triton) before further processing.

### Western blotting

Primary antibodies used were: Guinea pig anti-SMG (1:20000), rabbit anti-Smg (1:500), rat anti-Cup (1:10000), rabbit anti-Cup (1:1000), rabbit anti-TRAL (1:5000), rat anti-Tral (1:1000), rabbit anti-ME31B (1:5000), rabbit anti-Bel (1:2000) Guinea pig anti-SLMB (1:4000), rat anti-CG3295 (1:500), rat anti-eIF4E (1:1000). Mouse anti-Tubulin (1:20000) or mouse anti-Actin (1:1000) were used as loading controls.

For the developmental western blots shown in Figure 2 and Figure 2 – Figure supplement 1, embryos were collected, weighed, homogenized in SDS sample buffer with a pestle and boiled for 5 min. Proteins were resolved by SDS PAGE and transferred to a nitrocellulose membrane. Blots were blocked at room temperature with 1.5% cold water fish gelatin (Sigma) in TBS for 1 hour, and incubated with primary antibodies diluted in 1.5% gelatin in TBST (TBS plus 0.05% Tween 20) at 4°C overnight. Subsequently, blots were washed 5 × 6 minutes with TBST at room temperature and incubated with the appropriate fluorescently labeled secondary antibodies (1:15000, Li-COR) in TBST at room temperature for 2 hours. Blots were washed 5 × 6 minutes with TBST, imaged using an Odyssey CLx scanner and Image Studio Lite software (LI-COR) and band intensities were quantified using ImageJ.

For all other western blots, dechorionated embryos were counted, then lysed in SDS-PAGE sample buffer and boiled for 2 minutes. Proteins were resolved by SDS PAGE, and transferred to a PVDF membrane. Blots were blocked at room temperature with 2% non-fat milk in PBST for 30 minutes, and incubated with primary antibodies diluted in 2% non-fat milk in PBST at 4°C overnight. Subsequently, blots were washed 3× 10 minutes with PBST at room temperature and incubated with the appropriate HRP-conjugated secondary antibodies (1:5000, Jackson ImmunoResearch) in 2% non-fat milk in PBST at room temperature for 1 hour. Blots were washed 3× 15 minutes with PBST and developed using Immobilon Luminata Crescendo Western HRP substrate (Millipore), imaged using ImageLab (BioRad) and band intensities were quantified using ImageJ.

### MG132 treatment

1–2 h old embryos were permeabilized using a modification of a published method (Rand et al., 2010): Embryos were dechorionized for 30 s in 12% sodium hypochlorite. 45 ml of MBIM medium (Strecker et al., 1994) was mixed with 0.25 ml Triton X-100 and 4.5 ml (R)-(+)-limonene (Merck) and the dechorionated embryos were incubated with this mix for 30 seconds and then washed extensively with PBS followed by PBS/0.05% Tween 20. They were then incubated in MBIM containing 100 µM MG132, 0.05% DMSO or 0.05% DMSO for 3 hours at 25°C, washed in PBS, lysed in SDS-PAGE loading buffer and analyzed by Western Blot.

### Maternal RNAi knockdown

Females expressing Maternal-Gal4 driver were crossed to males expressing UAS-hairpin targeting each gene assayed from the TRiP library (see above). UAS-hairpin targeting mCherry was used as control knockdown. Adult F1s from these crosses were used for embryo collection for analysis. Maternal knockdown efficiency was assayed in 0–3h embryos by RT-qPCR. In the case of *slmb* RNAi, depletion of *slmb* mRNA was incomplete, and knockdown was further validated by Western blotting.

### FLAG-SMG Transgenes

For generation of transgenic flies expressing FLAG-SMG and FLAG-SMG767Δ999, the base vector used was the *smg5′UTR-BsiWI-smg3′UTR* (*SBS*) plasmid (Tadros et al., 2007)). A linker carrying a start codon, the FLAG/p53 epitope tags and AscI and PmeI restriction sites was inserted into the BsiWI site of *SBS*, between the *smg* UTRs. Genomic sequences of corresponding transgenic *smg* proteins were inserted between the AscI and PmeI sites. Coding sequence for amino acids 1-999 (for expressing full-length FLAG-SMG) and for amino acids 1-766 (for expressing truncated FLAG-SMG767Δ999 genomic transgene) were amplified from a *smg* genomic rescue construct (Dahanukar et al., 1999) using a 5′-primer with an AscI linker and a 3′-primer with PmeI. The ORFs were inserted between the AscI and PmeI sites in the SBS plasmid. *smg* genomic transgenes were then inserted into a pCaSpeR-4 cloning vector with an attB site (Markstein et al., 2008; Tadros et al., 2007). Transgenic *smg* constructs were integrated into an attP40 landing site on the second chromosome (2L:25C7) (Markstein et al., 2008) by Genetic Services (Cambridge, MA) using PhiC31, a site-specific integrase(Groth et al., 2004). The inserted transgenes were then crossed into a *smg*^*47*^ mutant background (Chen et al., 2014) for experiments.

### RT-qPCR

Total RNA was collected from dechorionated embryos in TRI Reagent (Sigma) following manufacturer protocol. 500ng of total RNA per sample was used to synthesize cDNA with Superscript IV reverse transcriptase kit (Invitrogen) using random hexamer primers. Primers specific to the transcripts assayed were designed using NCBI Primer-BLAST to cover all isoforms and span an exon-exon junction. Quantitative real-time PCR was performed using Sensifast SYBR PCR mix (Bioline) on a CFX384 Real-Time System (Bio-Rad). Expression of each gene was averaged across three technical replicates per biological replicate and normalized to *RpL32* control.

### Embryo developmental proteome

Each sample of ∼300 *y w* embryos was collected over a period of 1 hour at 22°C, then aged to the desired stage at the same temperature: (1) as early as possible – sample was not aged; The median time of this collection was defined as 0 min (2) Cycle 14 – 190 min; (3) germ-band extension – 330 min; (4) germ-band retraction – 630 min; and (5) trachea filling – 1290 min. The embryos were flash frozen in liquid nitrogen until lysis.

Samples were prepared as detailed previously (Gupta et al., 2018). The sample was pre-fractionated with a medium pH reverse chromatography. The fractions were analyzed via TMTc+ on an Orbitrap Fusion Lumos (Thermo Fisher). The LC-MS instrument was equipped with Easy nLC 1200 high pressure liquid chromatography (HPLC) pump. For each run, peptides were separated on a 100 μm inner diameter microcapillary column, packed first with approximately 0.5 cm of 5µm BEH C18 packing material (Waters) followed by 30 cm of 1.7µm BEH C18 (Waters). Separation was achieved by applying 6%-30% ACN gradient in 0.125% formic acid and 2% DMSO over 90 min at 350 nL/min at 60 °C. Electrospray ionization was enabled by applying a voltage of 2.6 kV through a microtee at the inlet of the microcapillary column. The Orbitrap Fusion Lumos was using the TMTc+ method (Sonnett et al., 2018b). The mass spectrometer was operated in data dependent mode with a survey scan ranging from 500-1400 m/z at resolution of 120k (200m/z). 10 most intense ions for MS2 fragmentation using the quadrupole. Only peptides of charge state 2+ were included. Dynamic exclusion range was set to 60 seconds with mass tolerance of 10ppm. Selected peptides were fragmented using 32% HCD collision energy, and the resultant MS2 spectrum was acquired using the Orbitrap with a resolution of 60k and 0.4 Th isolation window.

A suite of software tools (Huttlin et al., 2010; Sonnett et al., 2018a) was used to convert mass spectrometric data from the Thermo RAW file to the mzXML format, as well as to correct erroneous assignments of peptide ion charge state and monoisotopic m/z. Assignment of MS2 spectra was performed using the SEQUEST algorithm v.28 (rev. 12) by searching the data against the appropriate proteome reference dataset acquired from UniProt. This forward database component was followed by a decoy component which included all listed protein sequences in reversed order (Elias and Gygi, 2007). An MS2 spectral assignment false discovery rate of 0.5% was achieved by applying the target decoy database search strategy. Filtering was performed using a Linear Discriminant analysis with the following features: SEQUEST parameters XCorr and unique ΔXCorr, absolute peptide ion mass accuracy, peptide length, and charge state. Forward peptides within three standard deviation of the theoretical m/z of the precursor were used as positive training set. All reverse peptides were used as negative training set. Linear Discriminant scores were used to sort peptides with at least seven residues and to filter with the desired cutoff. Furthermore, we performed a filtering step on the protein level by the “picked” protein FDR approach(Savitski et al., 2015). Protein redundancy was removed by assigning peptides to the minimal number of proteins which can explain all observed peptides, with above described filtering criteria. TMTc+ data were analyzed as previously described (Sonnett et al., 2017). To correct for pipetting errors in the experiments, we normalized the signal assuming that the total protein amount remains constant (Tennessen et al., 2014).

### FLAG IP-MS

For FLAG-SMG IP/MS experiments, 0-3 hour embryos were collected from FLAG-SMG or non-transgenic *w*^*1118*^ flies as control. Dechorionated embryos were crushed in a minimal volume of lysis buffer (150 mM KCl, 20 mM HEPES-KOH pH 7.4, 1 mM MgCl2, 0.1% Triton X-100, supplemented with protease inhibitors and 1 mM DTT), cleared by centrifugation for 15 minutes at 4°C and 20000 x g, and stored at -80°C. Immediately prior to IPs, the lysate was diluted twofold with lysis buffer and cleared again by centrifugation. For each IP, 500μL of diluted lysate, with or without 0.35 µg/µL RNase A, was mixed with 20 µL of anti-FLAG M2 beads (Sigma; blocked with 5mg/mL BSA), and incubated for ∼3 hours at 4°C with end-over-end rotation. After incubation, beads were washed 4–5 times with lysis buffer, twice with lysis buffer lacking Triton X-100, then transferred to new tubes and washed twice with lysis buffer lacking Triton X-100. Bound proteins were eluted by tryptic digest: beads were resuspended in 200 µL of 50 mM ammonium bicarbonate, pH 8, with 2 µg of trypsin (Pierce), and incubated overnight at room temperature with end-over-end rotation. The digested supernatant was recovered, and beads were washed once with an additional 200 µL of 50 mM ammonium bicarbonate. The two eluates were pooled, dried by speed-vac, and analyzed by MS.

Samples were analyzed by liquid chromatography–tandem mass spectrometry (LC-MS/MS) using the Thermo Q-Exactive HF quadrupole-Orbitrap mass spectrometer (Thermo Scientific) and the methods previously described (Chiu et al., 2016; Jiang et al., 2015; Liu et al., 2014). Results were analyzed using the ProHits software package (Liu et al., 2010). For each protein detected, spectral counts were determined for peptides with iProphet probability > 0.95.

### GFP IP-MS

For GFP-tagged proteins, 0–2 hour old embryos were collected from flies expressing either GFP-SLMB, GFP-Muskelin or from non-transgenic Canton S flies as control. Embryos were dechorionated for 30 seconds in 12% sodium hypochlorite and lysed in 50 mM Tris pH 7.5, 150 mM sodium chloride, 1% NP-40, 0.5% sodium deoxycholate and protease inhibitor mix by homogenization with a pestle on ice. The extract was cleared by centrifugation (20000xg, 4°C), frozen in liquid nitrogen and stored at -80°C. The lysate was treated with RNase A (100 µg/ml) for 10 min at room temperature, cleared again by centrifugation (20000xg, 4°C) and diluted 1:5 with GFP wash buffer (50 mM Tris-HCl, pH 7.5, 150 mM NaCl, 0.5 mM EDTA). The diluted extract was incubated with GFP-Trap matrix (Chromotek, equilibrated in GFP wash buffer) for 1 h at 4°C. The matrix was washed 3 times with GFP wash buffer and resuspended in 8 M urea/0.4 M ammonium bicarbonate. Bound proteins were reduced with DTT (7.5 mM, 30 min at 50°C), alkylated with chloroacetamide (20 µl 100 mM per 120 µl reduced sample, 30 min at room temperature), digested with sequencing grade trypsin (Promega, 1:50 w/w trypsin to protein ratio) at 37°C overnight. Digests were stopped by addition of TFA, samples were concentrated by speed-vac and analyzed by LC/MS/MS using an U3000 RSCL nano-HPLC system coupled to an Orbitrap Q-Exactive Plus mass spectrometer with NanoFlex ionization source (all from Thermo Fisher Scientific). Samples were loaded on a trap column (PepMap RP-C18, 300 µm × 5 mm, 5 µm, 100 Å, Thermo Fisher Scientific) with 0.1 % TFA at a flow rate of 30 µl/min. After 15 min, peptides were eluted via an in-house packed separation column (self-pack PicoFrits, 75 µm × 50 cm, 15 µm tip, New Objective, packed with ReproSil-Pur 120 C18-AQ, 1.9 µm, Dr. Maisch GmbH). For peptide separation, a linear 180-min gradient was applied (3 % - 40 % eluent B; eluent A: 0.1 % formic acid in water, eluent B: 0.08 formic acid in acetonitrile) at a flow rate of 230 nl/min. Data were acquired using a data-dependent top10 strategy (one MS survey scan, followed by 10 MS/MS scans of the 10 most abundant signals). MS data (m/z range 375-1800) were recorded with R = 140,000 at m/z 200, MS/MS data (HCD, 28% normalized collision energy) with R = 17,500 at m/z 200. MS data were analyzed with MaxQuant 1.6.1.0 with label-free quantification (Cox and Mann, 2008). The modification parameter was set depending on the alkylation reagent used, iBAQ values were reported and the different runs of an experiment were matched if applicable.

### Determination of ubiquitination sites

For determination of ubiquitination sites, 1-3 hour old Canton S embryos were collected, dechorionated and lysed in 30 mM HEPES pH 7.6, 100 mM potassium acetate, 5 mM EDTA, 0.1% NP-40, 5 mM DTT, protease inhibitor mix, 50 µM MG132, 50 µM PR619 (100 µl per 100 mg embryos) by douncing. The extract was cleared by centrifugation (4× 7 min 20000 g), frozen in liquid nitrogen and stored at -80°C. 50µl of Protein A sepharose beads (GE Healthcare) were washed 4 x in IP buffer (16 mM HEPES pH 7.6, 50 mM potassium acetate, 1 mM EDTA, 0.1% NP-40) and incubated with 10µg of affinity-purified anti-Cup antibody overnight at 4°C, washed 3 times with IP buffer and incubated with 200 µl of extract (4°C, 3 hours). The beads were collected by centrifugation, washed 3 times with IP wash buffer (50 mM HEPES pH 7.6, 150 mM potassium acetate, 1 mM EDTA, 0.1% NP-40). During the last wash step, the matrix was transferred to new reaction tubes. The bound proteins were eluted twice with 300 µl 8 M urea, 0.4 M ammonium bicarbonate for 30 min at room temperature. The two eluates were pooled and acetone precipitated. The precipitates were resuspended in urea/ammonium bicarbonate, reduced with DTT (7.5 mM) for 30 min at 50°C, alkylated with 4-vinylpyridine (5 µl 100 mM per 35 µl reduced sample) for 30 min at room temperature, digested with sequencing grade trypsin (Promega, 1:50 w/w trypsin to protein ratio) at 37°C overnight. Digests were stopped by addition of TFA, samples were concentrated by speed-vac and analyzed by LC/MS/MS using the same instruments and settings as for GFP IP-MS; for analysis of ubiquitination, the modification K-GG was set in addition.

### Immunostaining

0–3 hour embryos were collected from F1 adults from maternal RNAi crosses, and dechorionated with 4.2% sodium hypochlorite for 2 minutes, fixed in 4% formaldehyde and heptane for 20 minutes, and devitellinized by the addition of methanol and vigorous shaking. Fixed embryos were rehydrated by PBSTx (PBS, 0.1% Triton-X 100) and blocked with 10% bovine serum albumin (BSA) in PBSTx. Embryos were incubated with guinea pig anti-SMG (1:2000 dilution, 1% BSA in PBSTx) rocking overnight at 4°C and subsequently washed 3 × 15 minutes while rocking at room temperature. Embryos were incubated in Cy3-conjugated donkey anti-guinea pig secondary antibody (1:300 dilution, 1% BSA in PBSTx; Jackson ImmunoResearch) for 1 hour rocking at room temperature, and washed 5 × 10 minutes with PBSTx. 0.001mg/ml DAPI (Sigma) was added to the second wash to label DNA. Embryo were mounted in 2.5% DABCO (Sigma), 70% glycerol in PBS. Images were collected using the Zeiss AxioSkop-2 MOT fluorescence microscope and the QCapture Suite PLUS acquisition software.

### SMG target RNA prediction

To computationally predict the formation of SMG recognition element (SRE) stem/loops (CNGGN_0-4_ loop sequence on a non-specific stem) within a transcript, we used a multi-step pipeline modified from (Chen et al., 2014). Each transcript was first scanned with RNAplfold (ViennaRNA package version 2.3.1) using the parameters -W=170 -L=120 -T=25 (Lange et al., 2012). Next, the transcript was scanned for CNGG motif (and variant motif) sites, and if the RNAplfold results indicated that the base immediately 5′ to the motif formed a base-pair interaction with one of the five nucleotides immediately 3′ to the motif with a probability > 0.01, then the motif was marked for further analysis. The probability of stem/loop formation at each site was then assessed with RNAsubopt using a 120nt sliding window that overlapped the candidate site (the first window beginning at 75nt upstream of the motif and extending 40nt downstream of the motif, which was shifted by one nucleotide in the 3′ direction per window 34 times for a total of 35 windows), for which 3000 structures were sampled per window. The empirical probability of stem/loop formation for each site was averaged across the 35 windows and expressed as a percentage to produce a score for each motif; scores from individual motifs were then summed across the entire to produce the final SRE score for the whole transcript. Zygotic targets not likely to be potential SMG targets had SRE scores < 5. All SMG-bound targets assayed had SRE scores > 10.

## ACKNOWLEDGEMENTS

We thank Olivia Rissland for communicating unpublished results prior to submission of her manuscript; Andrea Sinz for supporting the MS measurements at the University of Halle; Paul Schedl, Eric Wieschaus and Trudi Schüpbach for helpful advice and suggestions and Lillia Ryazanova for technical assistance at Princeton University; Luz Irina Calderon Villalobos and Michael Niemeyer for helpful discussions; and Thomas Hurd for a critical review of the manuscript. Antibodies were kindly provided by Akira Nakamura (rabbit anti-Cup, TRAL, ME31B) and Gregory Rogers (guinea pig anti-SLMB). W.X.C. was supported in part by an Ontario Graduate Scholarship and University of Toronto Open Scholarships. This research was supported by the following funding agencies: Canadian Institutes of Health Research (PJT-159702 to H.D.L.), Natural Sciences and Engineering Research Council of Canada (to H.D.L. and to C.A.S.), Deutsche Forschungsgemeinschaft (WA 548/17-1 within SPP 1935 to E.W.), National Institute of Health (R35GM128813 to M.W. and T32GM007388 to E.Y.).

## CONTRIBUTIONS

The project was designed by H.D.L. and E.W. The unfertilized egg time-course, RNAi experiments, analyses of FLAG-SMG and FLAG-SMG767Δ999 time-course westerns, and RT-qPCR experiments to assess mRNAs in FLAG-SMG and FLAG-SMG767Δ999 were carried out by W.X.C. under the supervision of H.D.L. ; the RBP western time-course, MG132 experiments and GFP-Muskelin/CG3295 westerns were carried out by S.K. under the supervision of E.W.; TMTc+ mass spectrometry was carried out by M.G. and E. Y. under the supervision of M.W.; FLAG-SMG IP-MS was carried out by W.X.C. and S.L. under the supervision of H.D.L and S.A., respectively; GFP-Muskelin, GFP-SLMB and Cup IP-MS was carried out by S.K., C.R. and C.I. under the supervision of E.W.; F.P. generated the anti-eIF4E and anti-CG3295 antibodies under the supervision of E.W.; T.C.H.L. wrote the script to calculate SRE scores under supervision of H.D.L.; N.U.S. constructed the transgenes to express FLAG-SMG and FLAG-SMG767Δ999 under the supervision of H.D.L.; M.H.K.C. showed that FLAG-SMG767Δ999 persists under the supervision of C.A.S.. W.X.C. wrote the first draft of the manuscript, which was revised by H.D.L. with major input from E.W. and M.W., as well as input from the other co-authors.

---

**Figure 2 – Figure supplement 1.**
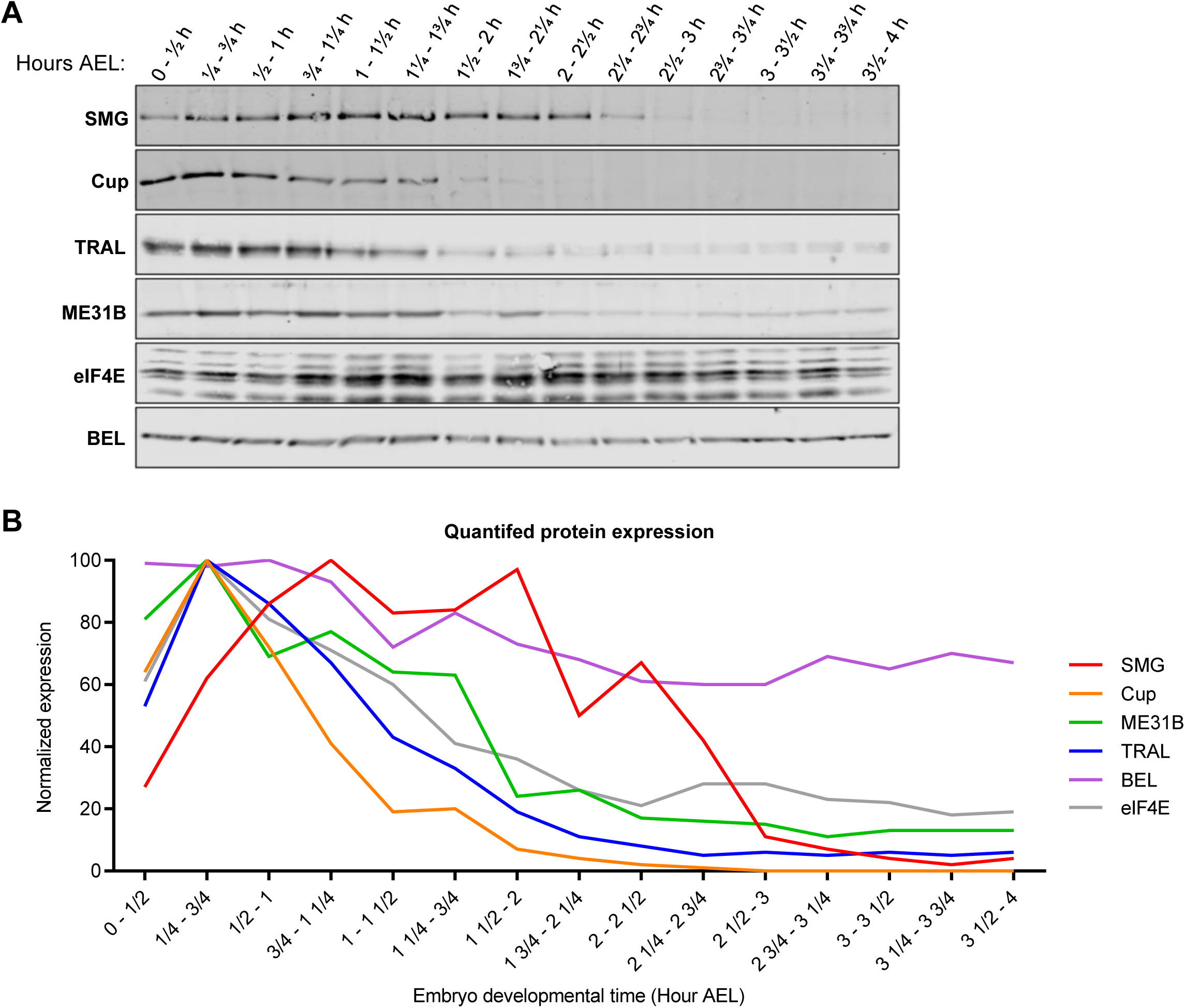
RBP expression is differentially regulated during the MZT. **A.** Biological replicate of the developmental western blot shown in **Figure 2A**, assayed over the first four hours of embryogenesis after egg-lay (AEL). **B.** Quantification of protein expression across 2 biological replicates and at least 2 technical replicates (with the exception that only one sample was quantified for eIF4E). Each protein was normalized to tubulin (see Figure 2, not shown here), and its peak expression during the time course was set to 100.

**Figure 4 – Figure supplement 1.**
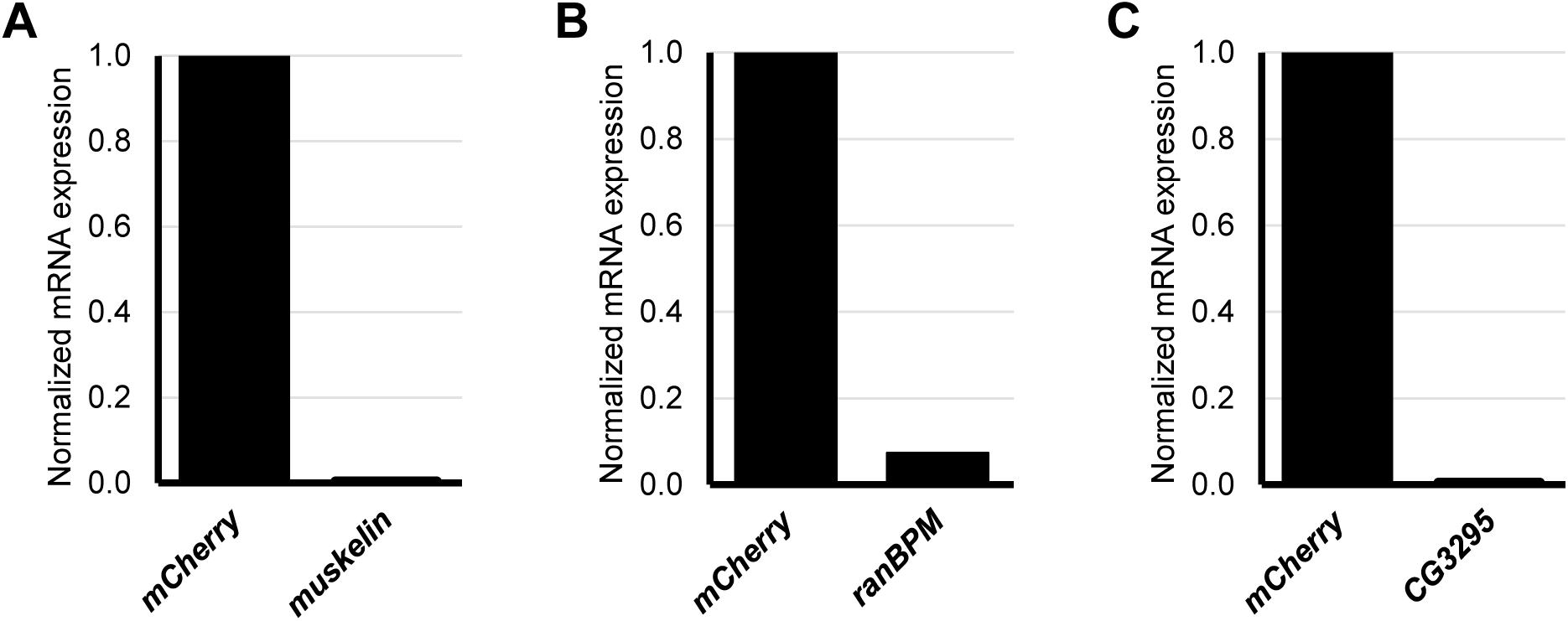
Validation of maternal RNAi knockdown of CTLH complex members. RT-qPCR quantification of target mRNA expression in 0-3h embryos. **A.** *muskelin*, **B.** *ranBPM* and **C.** *CG3295* mRNAs were depleted by > 90% relative to *mCherry* control RNAi in their respective maternal knockdowns.

**Figure 4 – Figure supplement 2.**
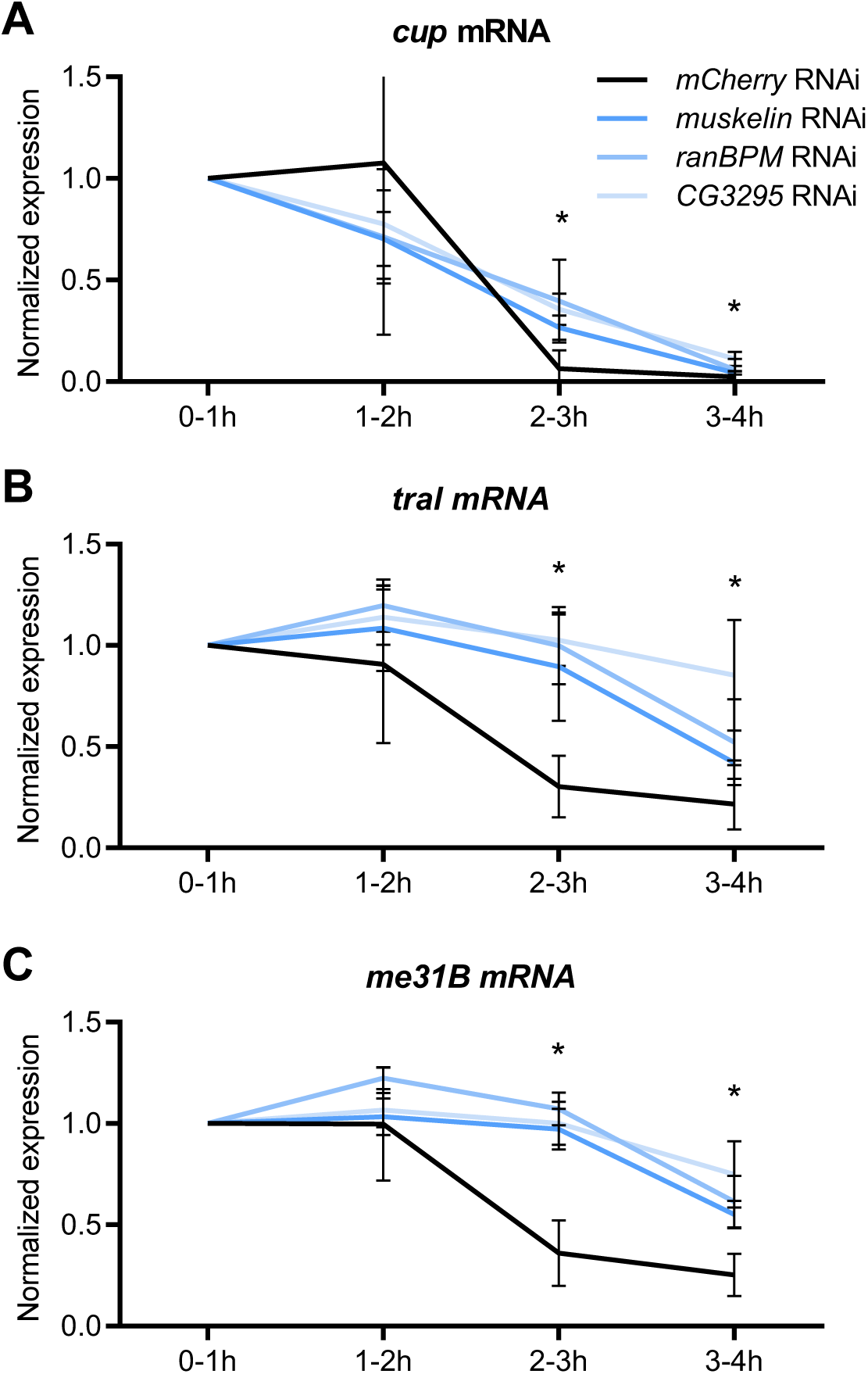
Knockdown of the CTLH complex results in delayed degradation of mRNAs. RT-qPCR quantification of mRNA expression in embryos assayed in **Figure 4A-D**. Knockdown of the CTLH complex resulted in delayed degradation of *cup* (**A**), *tral* (**B**) and *me31B* (**C**) mRNA relative to control knockdown. *tral and me31b* remained partially stabilized at 3-4h after egg-lay. \**P* < *0.05, n.s. =* not significant, n = 3, error bars = SD, Student’s *t*-test.

**Figure 4 – Figure supplement 3.**
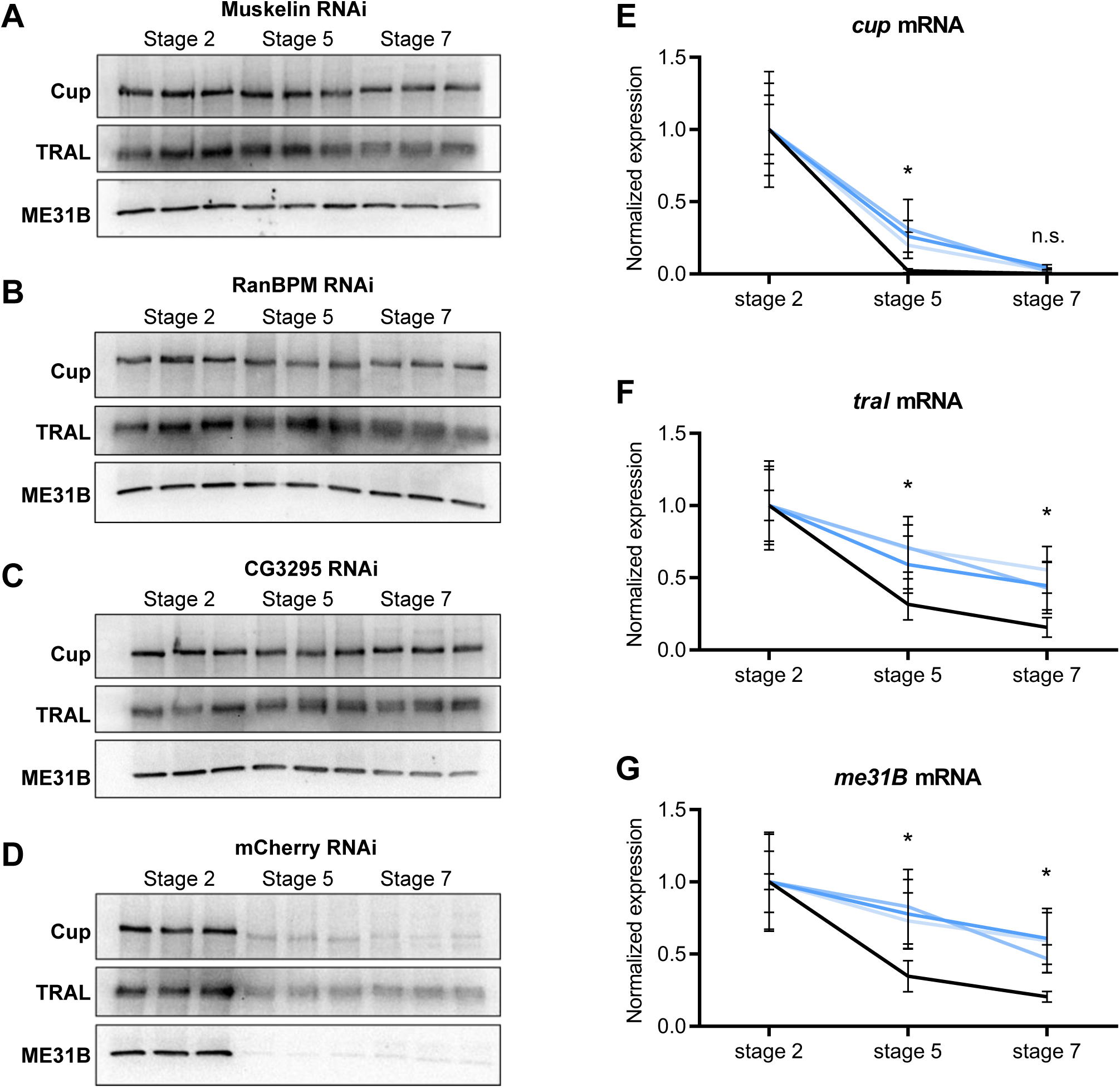
Knockdown of the CTLH complex stabilizes Cup in developing embryos. **A-D.** Western blots of embryos picked at Stage 2, Stage 5 and Stage 7 (n = 3 biological replicates for each stage). Maternal knockdown of *muskelin* (**A**), *ranBPM* (**B**) and *CG3295* (**C**) resulted in stabilization of Cup, TRAL and ME31B through embryo Stage 7, whereas all three RBPs were depleted in the embryo by Stage 5 in the control knockdown (**D**)**. E-G.** RT-qPCR quantification of mRNA expression in embryos assayed in A-D. Picking developmentally stage-matched embryos partially rescued the delay in mRNA degradation in CTLH maternal knockdown embryos. *cup* (**E**) was cleared to comparable levels as in control *mCherry* RNAi by stage 7. *tral* (**F**) and *me31B* (**G**) remained partialized stabilized. \**P* < *0.05, n.s. =* not significant, n = 3, error bars = SD, Student’s *t*-test.

**Figure 5 – Figure supplement 1.**
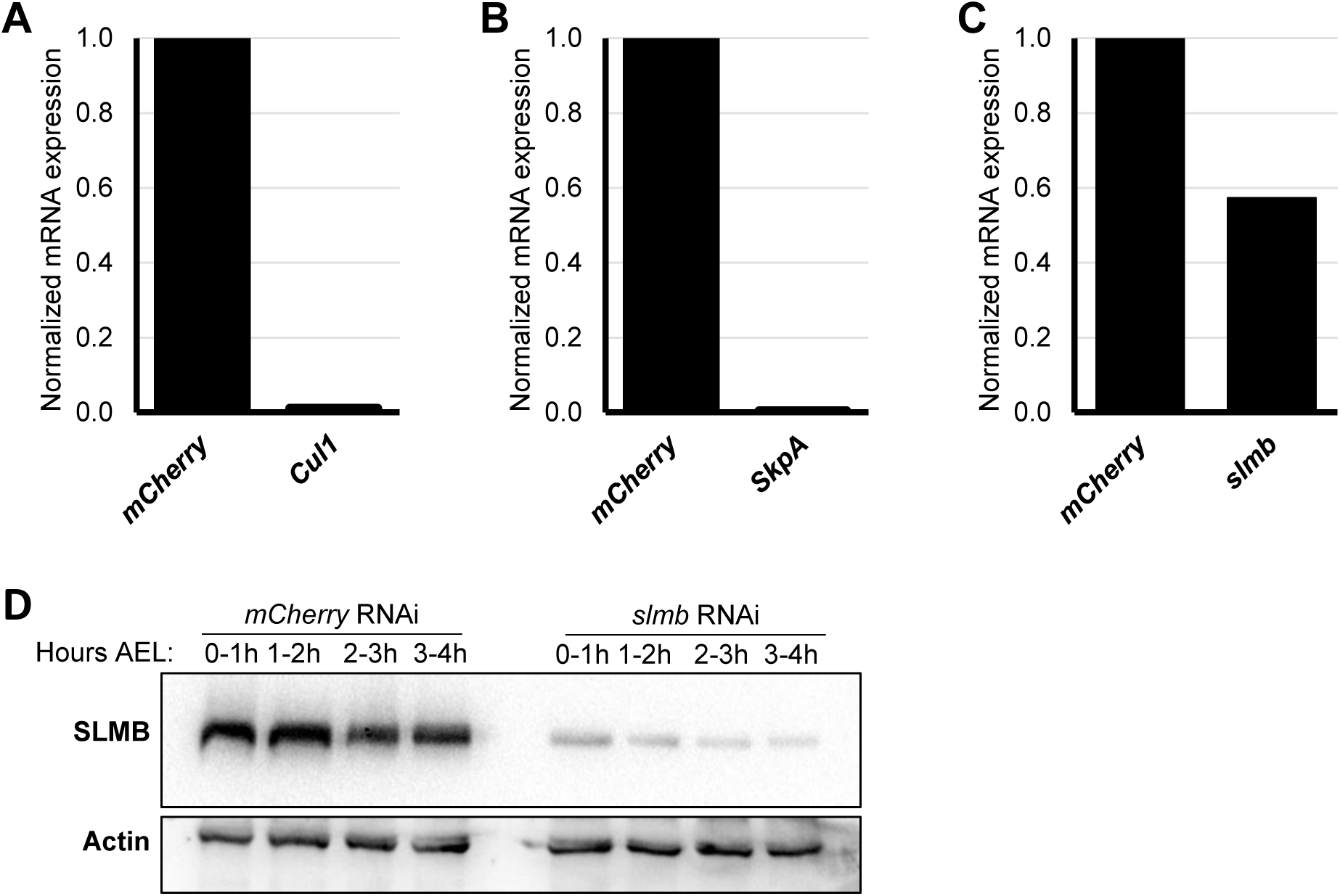
Validation of maternal RNAi knockdown of SCF complex members. RT-qPCR quantification of target mRNA expression in 0-3h embryos. **A.** *cul1* and **B.** *skpA* mRNAs were depleted by >98% relative to *mCherry* control RNAi in their respective maternal knockdowns. **C.** Maternal knockdown of *slmb* achieved a 43% reduction at the RNA level. **D.** Western blot of SLMB protein expression in *mCherry* control RNAi and *slmb* RNAi showed ∼90% depletion of SLMB protein expression over the first 4 hours of embryogenesis resulting from the maternal *slmb* knockdown. Note: The same blot shown here is cropped and used in **Figure 6**.

**Figure 5 – Figure supplement 2.**
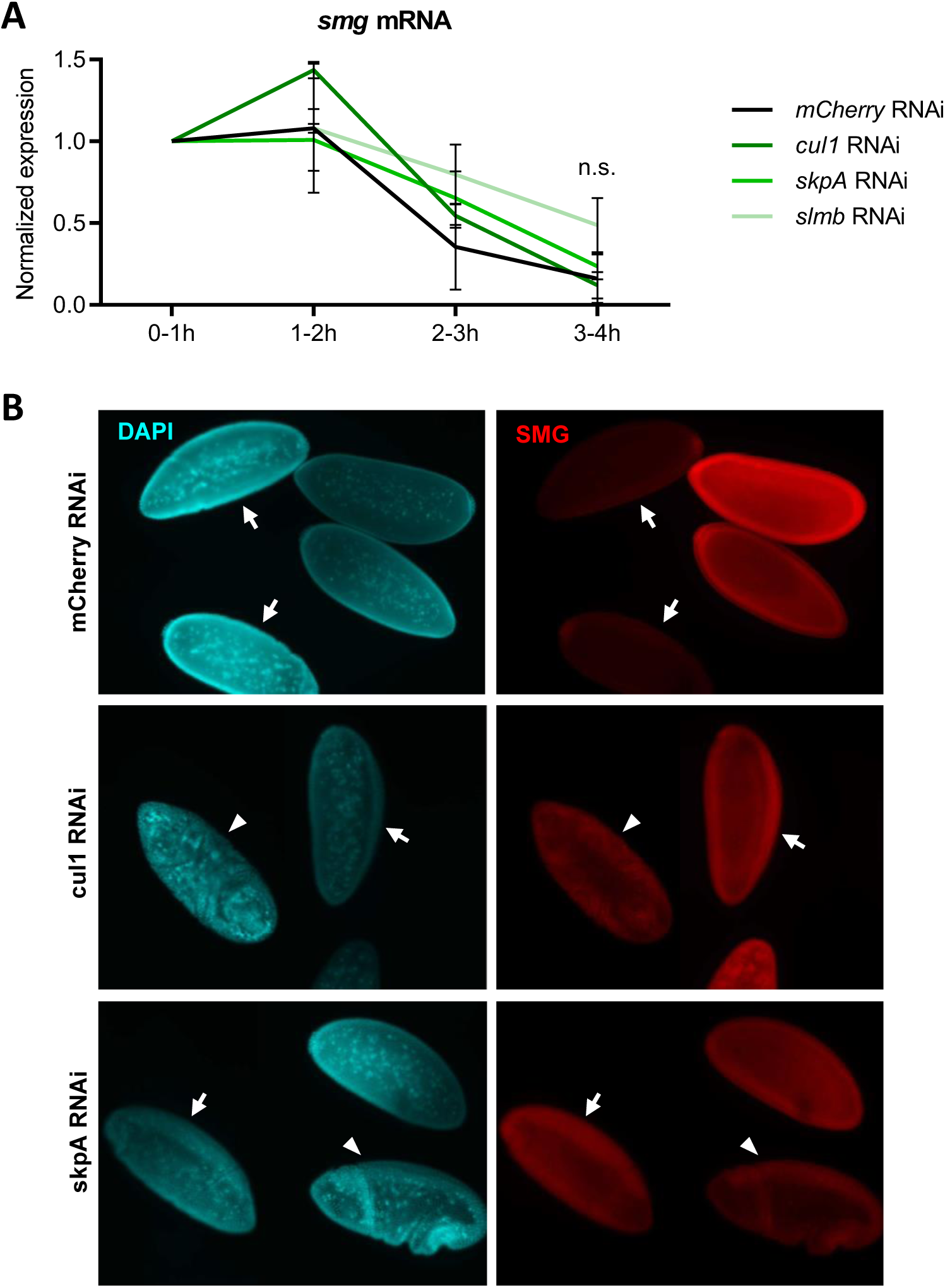
Knockdown of the SCF complex stabilizes SMG protein independent of RNA degradation and embryo development. **A.** RT-qPCR quantification of *smg* mRNA expression in embryos assayed in **Figure 5D**. Knockdown of the SCF complex had no significant effect on *smg* mRNA degradation relative to control knockdown. *n.s. =* not significant, n = 3, error bars = SD, Student’s *t*-test. **B.** Immunofluorescence of nuclei (DAPI, blue) and SMG (red) in maternal knockdown embryos. In *mCherry* control knockdown, SMG was depleted from the blastoderm embryo by the onset of gastrulation (arrows). Knockdown of *cul1* or *skpA* resulted in ubiquitous persistence of SMG protein in comparable stages, as well as older gastrulating embryos (arrowheads).

